# Mechanism of activation and regulation of Deubiquitinase activity in MINDY1 and MINDY2

**DOI:** 10.1101/2021.01.27.428544

**Authors:** Syed Arif Abdul Rehman, Lee A. Armstrong, Sven M. Lange, Yosua Adi Kristariyanto, Tobias W. Grawert, Axel Knebel, Dmitri I. Svergun, Yogesh Kulathu

## Abstract

Of the eight distinct polyubiquitin chains that can be assembled, K48-linked ubiquitin is the most well-understood linkage and modification of proteins with K48 chains targets the modified protein for degradation. By removing ubiquitin from substrates or trimming ubiquitin chains, deubiquitinases (DUBs) can modulate the outcome of ubiquitylation. MINDY1 and MINDY2 are members of the MINDY family of DUBs that have exquisite specificity for cleaving K48-linked polyubiquitin. Being recently discovered DUBs, we have a poor understanding of their catalytic mechanism. By analysing crystal structures of MINDY1 alone and in complex with monoubiquitin or K48-linked ubiquitin chains, we here reveal how substrate interaction relieves autoinhibition and activates the DUB. Further, our analyses reveal a non-canonical catalytic triad composed of Cys-His-Thr and explain how these DUBs sense both ubiquitin chain length and linkage type to trim K48-linked ubiquitin chains. Our findings highlight the multiple layers of regulation modulating DUB activity in MINDY1 and MINDY2.

**Synopsis:**

- Structure of MINDY1 in complex with K48-linked diUb reveals how K48-linked polyUb is recognized and cleaved
- The Cys loop mediates autoinhibition of the DUB and substrate binding at the S1 and S1’ sites relieves autoinhibition and activates the enzyme for catalysis
- MINDY1 uses a non-canonical catalytic triad composed of Cys-His-Thr
- MINDY1 has five ubiquitin binding sites within its catalytic domain and switches from exo to endo cleavage in a ubiquitin chain length-dependent manner

## INTRODUCTION

Nearly all aspects of eukaryotic cell biology are influenced by the posttranslational modification (PTM) of proteins with ubiquitin (Ub). Typically, Ub is tagged onto a substrate protein by the formation of an isopeptide bond between its C-terminal carboxyl group and the ε-amine group of a lysine residue on the substrate. This primary Ub can be extended where one of its seven lysine residues (K6, K11, K27, K29, K33, K48, K63) or its N-terminal methionine (M1) can serve as an attachment point for another ubiquitin to result in the formation of polyUb chains (Komander & Rape, 2012). Importantly, the linkage type of the ubiquitin chain can determine the fate of the modified protein and the functional outcome of ubiquitylation. The most common function of ubiquitylation is mediated by K48-linked ubiquitin chains, which target the modified protein for destruction by the proteasome (Hershko & Ciechanover, 1998). K48-linked ubiquitylation is therefore the most prevalent modification in cells as it is essential to degrade proteins in a timely manner to maintain proteostasis, the failure of which is the underlying cause for many diseases including neurodegenerative diseases (Dikic, 2017).

Given their wide-ranging effects on cell signalling and eukaryotic biology, it is essential for cells to tightly regulate ubiquitylation. This regulation is largely mediated by dedicated ubiquitin proteases called deubiquitinases (DUBs) which can remove ubiquitin from the substrate or trim polyubiquitin chains (Clague *et al*, 2019; Leznicki & Kulathu, 2017). There are around 100 DUBs known so far in humans which are broadly classified into seven different families based on their structural fold and catalytic mechanism (Kwasna *et al*, 2018; Abdul Rehman *et al*, 2016; Ronau *et al*, 2016; Clague *et al*, 2019) Of these seven families, six are cysteine proteases while one family is made up of metalloproteases. The majority of DUBs belonging to the USP family show no linkage preference and can hydrolyze all polyubiquitin chain types. In contrast, DUBs belonging to the OTU, MINDY and ZUP1 classes exhibit exquisite linkage selectivity and cleave only specific linkage types (Faesen *et al*, 2011; Mevissen *et al*, 2013; Abdul Rehman *et al*, 2016; Kwasna *et al*, 2018).

The mode of ubiquitin recognition by the DUB determines whether it is linkage specific. For instance, USP family DUBs have a large S1 ubiquitin binding site and their activity depends on distal ubiquitin binding. On the other hand some DUBs rely on additional proximal ubiquitin interactions at the S1’ site to orient a specific lysine or the N-ter methionine towards the catalytic site, thus making them selective at cleaving specific linkages (Clague *et al*, 2019). The S1’ site on a DUB may be present within the catalytic domain or provided by a ubiquitin binding domain (UBD) in the DUB (Keusekotten *et al*, 2013; Mevissen *et al*, 2013; Clague *et al*, 2019). A key feature of these ubiquitin interactions is to stabilize the linkage between Ub moieties in a Ub chain within the active site for cleavage. Ubiquitin chains of different linkage types adopt distinct conformations. K33-, K63- and M1-linked chains can exist in an open extended conformation with accessible I44 binding patches that can be recognized by DUBs (Kristariyanto *et al*, 2015). Other linkage types such as K6-, K11- and K48-linked chains adopt compact conformations and must undergo significant conformational changes to be recognized by a DUB (Mevissen *et al*, 2016; Gersch *et al*, 2017; Sato *et al*, 2017; Ye *et al*, 2012). With K48-linked chains, the I44 patches are tightly buried at the interface and so we do not know how this linkage type is recognized by DUBs as there are no structures of DUBs bound to K48-linked chains available.

DUBs also process ubiquitin chains in different ways to modulate ubiquitylation, and the mode of chain cleavage is a factor that can influence the duration of the Ub signal. For instance, DUBs can cleave from one end of the chain (exo-DUB) to trim the ubiquitin chain (Leznicki & Kulathu, 2017). Alternatively, DUBs can hydrolyse within ubiquitin chains in a mode of endo cleavage that can rapidly terminate a ubiquitin signal and is characteristic of DUBs such as CYLD and A20 that regulate ubiquitin signalling (Komander *et al*, 2008; Lin *et al*, 2008). Enzymes are subject to multiple layers of regulation that ensure activity at a precise time and location. Similarly, it is essential that the activity of DUBs is modulated. Several DUBs exist in an autoinhibited conformation typified by a misaligned catalytic triad or occluded substrate-binding site, and depend on post-translational modifications, allosteric interactions or substrate interactions for activation (Sahtoe & Sixma, 2015).

We recently reported the discovery of the MINDY (MIU containing novel DUB family) enzymes as a novel class of cysteine protease DUBs. DUBs of the MINDY family are evolutionarily conserved and are all remarkably specific at cleaving K48-linked chains (Abdul Rehman *et al*, 2016). MINDY1 and MINDY2 show high sequence similarity and similar domain architectures. We found that MINDY1 is an exo-DUB with a preference for cleaving long K48-linked polyUb chains. Further, MINDY1 exists in an inhibited conformation and the identity of the catalytic triad is unclear. Hence, the catalytic mechanism, and how MINDY1 gets activated and specifically cleaves K48-linked ubiquitin chains in this unique fashion, is unknown. Moreover, K48 chains exist in a closed compact conformation raising the question of how these compact structures are recognized by MINDY1. In fact, to our knowledge there are no structures available of any DUB in complex with K48 chains with the isopeptide bond positioned across the catalytic site. To address these questions, we here determined the crystal structures of MINDY1 and MINDY2 in complex with K48-linked diUb (K48-Ub_2_). Our structural analyses coupled with mutational studies reveal the mechanism of autoinhibition and activation of MINDY1 and MINDY2.

## RESULTS

### Structure of MINDY1 in complex with K48-linked diUb

The minimal catalytic domain of MINDY1 (MINDY1^cat^) contains all the specificity determinants to selectively cleave K48-linked chains (Abdul Rehman *et al*, 2016). To understand the structural basis for the specific cleavage of K48-linked chains by MINDY1^cat^, we determined the crystal structure of a catalytically dead (C137A) mutant of MINDY1^cat^ bound to K48-linked diUb (MINDY1:K48-Ub_2_) to 2.2 Å resolution (**Table 1**). The structure was solved by molecular replacement with MINDY1^cat^ (PDB 5JKN) (Abdul Rehman *et al*, 2016) and ubiquitin (PDB 1UBQ) (Vijay-kumar *et al*, 1987) as search models. In the crystal, MINDY1 forms a stoichiometric complex with K48-Ub_2_ with one complex in the asymmetric unit (ASU) (**Fig 1A**).

**Table 1:** Data collection and refinement statistics.

|  | **MINDY1:K48-Ub2** | **MINDY1<sup>P138A</sup>** | **MINDY1<sup>P138A</sup>:K48-Ub<sub>2</sub>** |
| --- | --- | --- | --- |
| **Data Collection** |  |  |  |
| Beamline | ID23-1, ESRF | ID23-1, ESRF | ID30B, ESRF |
| Wavelength (Å) | 0.939274 | 0.97625 | 0.99187 |
| Space group | P4<sub>1</sub>22 | P4<sub>1</sub>22 | P4<sub>1</sub>22 |
| Total reflections | 212300 (19256) | 86341 (20676) | 213260 (18229) |
| a, b, c (Å) | 73.86, 73.86, 202.87 | 98.65, 98.65,<br>166.07 | 74.03, 74.03,<br>202.85 |
| α, β, γ (°) | 90.00, 90.00, 90.00 | 90.00, 90.00, 90.00 | 90.00, 90.00,<br>90.00 |
| Resolution (Å) | 41.81-2.16 | 48.28-3.59 | 41.84-2.18 |
| R<sub>merge</sub> | 0.082 (0.923) | 0.089 (0.977) | 0.066 (0.902) |
| I/σ(I) | 13.9 (4.1) | 12.5 (2.3) | 14.2 (2.0) |
| Completeness (%) | 99.7 (99.2) | 99.9 (99.9) | 100.0 (100.0) |
| Multiplicity | 6.8 (7.3) | 8.5 (8.8) | 7.0 (7.1) |
| CC1/2 | 0.999 (0.889) | 0.999 (0.905) | 0.999 (0.819) |
| **Refinement** |  |  |  |
| Resolution (Å) | 41.37-2.16 | 48.00-3.59 | 41.84-2.180 |
| R<sub>work</sub>/ R<sub>free</sub> | 0.207/0.247 | 0.212/0.258 | 0.212/0.246 |
| **No. of Atoms** |  |  |  |
| Protein | 3451 | 1946 | 3571 |
| Ligand | 104 | N/A | 6 |
| Water | 5 | N/A | 57 |
| **B-Factors (Å<sup>2</sup>)** |  |  |  |
| Protein | 60.41 | 191.19 | 75.24 |
| Ligand | 58.20 | N/A | 82.71 |
| Water | 48.13 | N/A | 49.34 |
| **RMSDs** |  |  |  |
| Bond length (Å) | 0.009 | 0.006 | 0.006 |
| Bond angles (°) | 1.608 | 1.670 | 1.036 |
| **Ramachandran Plot (%)** |  |  |  |
| Favored region | 97 | 83 | 98 |
| Allowed region | 3 | 17 | 2 |
| Outlier region | 0 | 0 | 0 |
| PDB ID | **6TUV** | **6Z90** | **6TXB** |

DLS, Diamond Light Source; ESRF, European Synchrotron Radiation Facility; RMSD, root-mean-square deviation. Values for the highest-resolution shell are shown in parentheses

**Figure 1:**
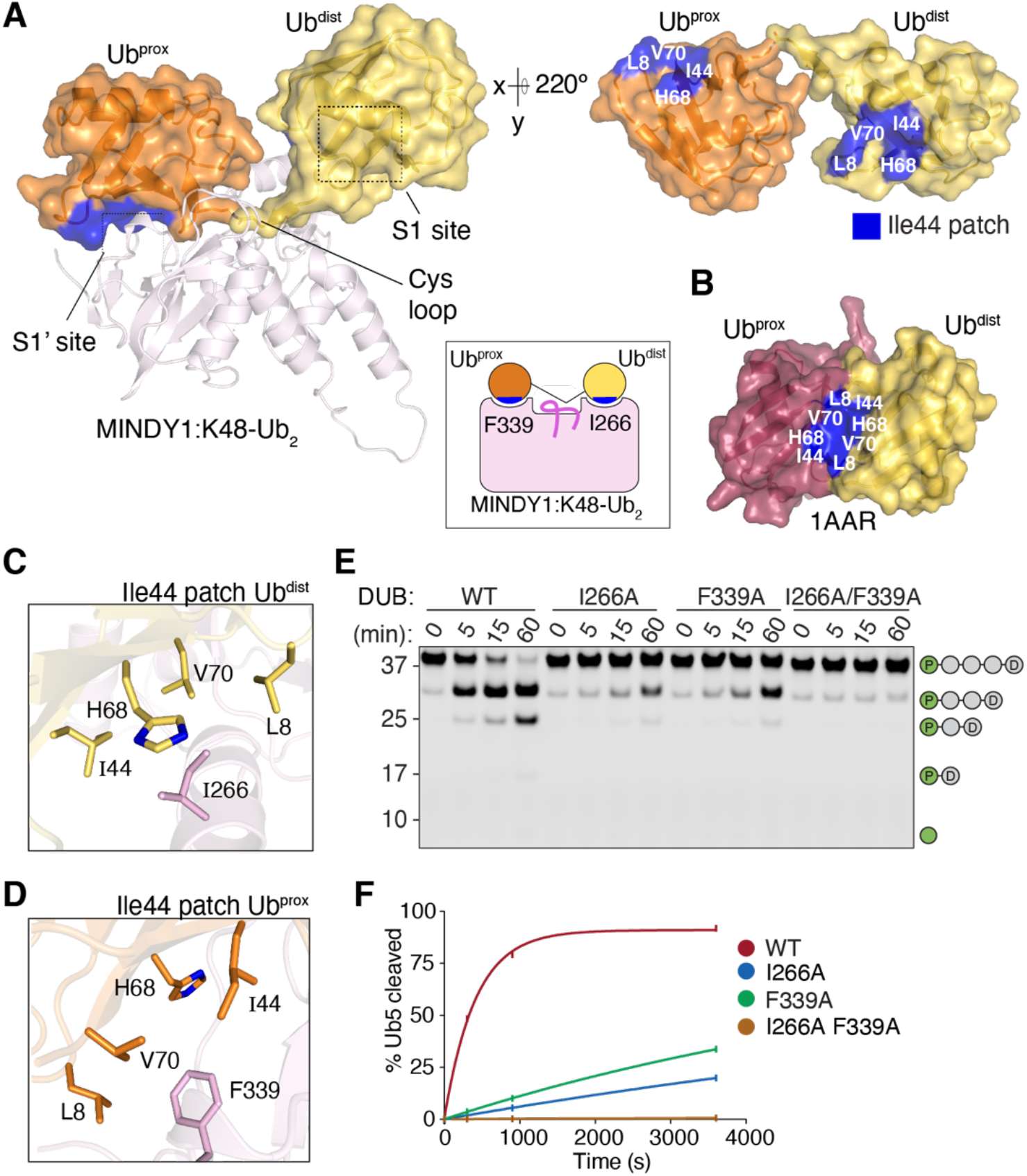
Crystal structures of MINDY1 in complex with K48 linked di-ubiquitin. **A**) The MINDY1:K48-Ub2 complex crystal structure is shown with MINDY1 in cartoon (light pink). Ub molecules are depicted with transparent surfaces (tv-orange:Ubprox and yelloworange: Ubdist). I44 patches on Ub are coloured blue and an alternate view of the bound diUb rotated by 220º along the x-axis is shown on the right side. Schematic representation of MINDY1:K48-Ub2 complex (inset). **B**) Surface representation of the closed conformation of K48-Ub2 (PDB ID:1AAR) with I44 patches highlighted in blue **C-D**) Close-up views of the key residues on the MINDY1 S1 and S1’ sites and their interactions with the I44 patches on Ubdist and Ubprox. **E**) DUB assay monitoring cleavage of K48-linked pentaUb, in which Ubprox is fluorescently labelled, by MINDY1 and indicated mutants. **F**) Quantification of pentaUb hydrolysis shown in panel D. The percentage of the total intensities of Ub4, Ub3, Ub2, and Ub1 formed is shown on the y-axis.

MINDY2 displays close sequence similarity to MINDY1 and is also selective at cleaving K48 chains. Further, the minimal catalytic domain of MINDY2 (MINDY2^cat^) is also specific for cleaving K48 chains (**Fig S1A,B**). To understand if similar mechanisms regulate specificity and activity in MINDY2, we determined the crystal structure of MINDY2^cat^ alone and catalytically dead MINDY2^cat^ (C266A) in complex with K48-Ub_2_ (**Table 1**). The crystal structure of MINDY2^cat^ closely resembles that of MINDY1^cat^ (RMSD 1.05Å) and binds to K48 chains in a similar way (**Fig S1C**). Overall, MINDY1 and MINDY2 use similar mechanisms and for the sake of simplicity, we will focus on MINDY1 in our description highlighting any differences observed with MINDY2.

The crystal structure of MINDY1: K48-Ub_2_ complex reveals how MINDY1^cat^ has extensive interactions with both the proximal Ub (Ub^prox^) and distal Ub (Ub^dist^) to position the scissile bond across the catalytic site (**Fig 1A, S1D**). We observe electron density for both Ub^prox^ and Ub^dist^ (**Fig S1E,F**). While the C terminal G76 of the distal Ub points towards K48 of the proximal Ub, electron density for the isopeptide bond itself is not visible (**Fig S1E**). However, electron density for the isopeptide bond is visible in the MINDY2: K48-Ub_2_ complex (**Fig S1F**).

The distal Ub binds to the S1 site in MINDY1 with a buried surface area of only ~750 Å^2^. Of note, this S1 binding interface is much smaller than those of other DUBs such as HAUSP/USP7 (1700 Å^2^), USP21 (1700 Å^2^) or USP2 (1900 Å^2^), which rely more on Ub^dist^ binding for activity (Ye *et al*, 2011; Renatus *et al*, 2006; Hu *et al*, 2002). In the crystal structure of MINDY1:Ub^Prg^ complex, the ubiquitin occupying the S1 site exists in two alternate conformations suggestive of weak binding. One of these conformers, conformer A, corresponds to Ub^dist^ in the K48-Ub_2_ complex (**Fig S2C**). This suggests that Ub^prox^ interactions with MINDY1^cat^ stabilizes Ub^dist^ binding. Indeed, the proximal Ub has a slightly larger binding interface and binds with a buried surface area of ~965 Å^2^. Furthermore, the residues involved in proximal and distal Ub recognition in MINDY1 are conserved in evolution (**Fig S2A,B**), suggesting that simultaneous binding of both Ub^dist^ and Ub^prox^ is essential to properly position the scissile bond for catalysis.

In both the MINDY1:K48-Ub_2_ and MINDY2:K48-Ub_2_ complexes, the K48-linked chain adopts an extended conformation that lacks interchain Ub-Ub interactions (**Fig 1A, S1D**). This is in contrast to the compact conformation observed for K48-linked chains in isolation in which the I44 patches of both Ub^dist^ and Ub^prox^ form a hydrophobic interface (**Fig 1B**). In this extended conformation, the I44 patches of both Ub^prox^ and Ub^dist^ are now engaged in interactions with the catalytic domain (**Fig 1A**). The I44 patch on Ub^dist^ primarily contacts I266 on MINDY1 (**Fig 1C**). Additional interactions with Ub^dist^ are mediated by V210, W240, Y258 and F315 in MINDY1, which form a binding pocket for L73 of Ub^dist^, and polar interactions between D209 and E263 with R74 and R72 of Ub^dist^ further contribute to binding (**Fig S2D,E**). Similar to other DUBs, Ub^dist^ L73 interaction is important and mutating the L73 binding pocket on MINDY1 impairs DUB activity (Békés *et al*, 2016; Abdul Rehman *et al*, 2016). The I44 patch of Ub^prox^ mainly contacts F339 on MINDY1 (**Fig 1D**). Additional cation-π interactions between Ub^prox^ F45 and MINDY1 R316, hydrogen bonds between the side chains of N317 and Ub^prox^ N60 and the backbone of Y59, and ionic interactions between Ub^prox^ R42 and MINDY1 D336 establish Ub^prox^ binding at the S1’ site (**Fig S2D-F**). In summary, hydrophobic interactions with the I44 patch on both ubiquitin moieties together with additional ionic interactions and hydrogen bonds help to precisely position the scissile bond of the K48 chain across the active site groove of MINDY1.

Monitoring the cleavage of fluorescently-labelled K48-Ub5 by the MINDY1 mutants I266A (S1 site) and F339A (S1’ site), which disrupt binding of Ub^dist^ and Ub^prox^ respectively results in reduced activity relative to WT, whilst a double mutant (I266A/F339A) completely abolishes DUB activity (**Fig 1E,F and S1G**). Thus, simultaneous engagement of the I44 patches of both Ub^prox^ and Ub^dist^ is essential for catalysis as disrupting either interaction hinders the ability of MINDY1 and MINDY2 to cleave K48 chains. Comparison of all available structures of DUBs in complex with diUb bound at the S1 and S1’ sites reveals that this mode of symmetric binding where the I44 patches of both Ub^prox^ and Ub^dist^ are engaged with the DUB is unique to MINDY1/2 (**Fig 1A** inset, **Table S1**). K48-linked chains exist in a dynamic equilibrium between closed and open conformations (Ye *et al*, 2012). Interestingly, the extended conformation adopted by K48-linked diUb when bound to the catalytic domains of MINDY1 and MINDY2 is distinct from all previously reported conformations of K48-linked chains (**Fig S2G**). Despite K48-linked polyUb being the first described and most well understood of all ubiquitin linkage types, to our knowledge these structures of MINDY1/2 represent the first crystal structures of any DUB in complex with K48-linked chains.

### Cys loop regulates DUB activity

In MINDY1 and MINDY2, a flexible loop (T130 to P138 in MINDY1 and T259 to P267 in MINDY2) connects β2 to α1, which we term the Cys loop since the catalytic cysteine sits at its base. This Cys loop occludes the catalytic centre and would sterically occlude positioning of the scissile bond across the active site (**Fig 2A**). Comparing the structures of MINDY1: K48-Ub_2_ complex with MINDY1^apo^ does not show any large-scale conformational changes induced within the catalytic domain upon ubiquitin binding (RMSD 1 Å over 244 aligned Cα atoms) (**Fig S3A**). The only significant change is in the Cys loop that is remodelled, during which several hydrogen bonds are broken accompanied by the formation of new bonds (**Fig 2B, S3B-D**). This Cys loop remodelling is observed in both MINDY1 and MINDY2 (**Fig S3G**) and when remodelled, the Cys loop no longer impedes ubiquitin binding. In the apo structure, in addition to the obstructing Cys loop, the catalytic residues are misaligned in an unproductive conformation (Abdul Rehman *et al*, 2016). Hence the apo state of the enzyme corresponds to an inactive or autoinhibited conformation and the DUB transitions to an active state when in complex with K48-linked diUb. We hypothesize that the dynamics of the Cys loop is important for the activation of MINDY1 and MINDY2.

**Figure 2.**
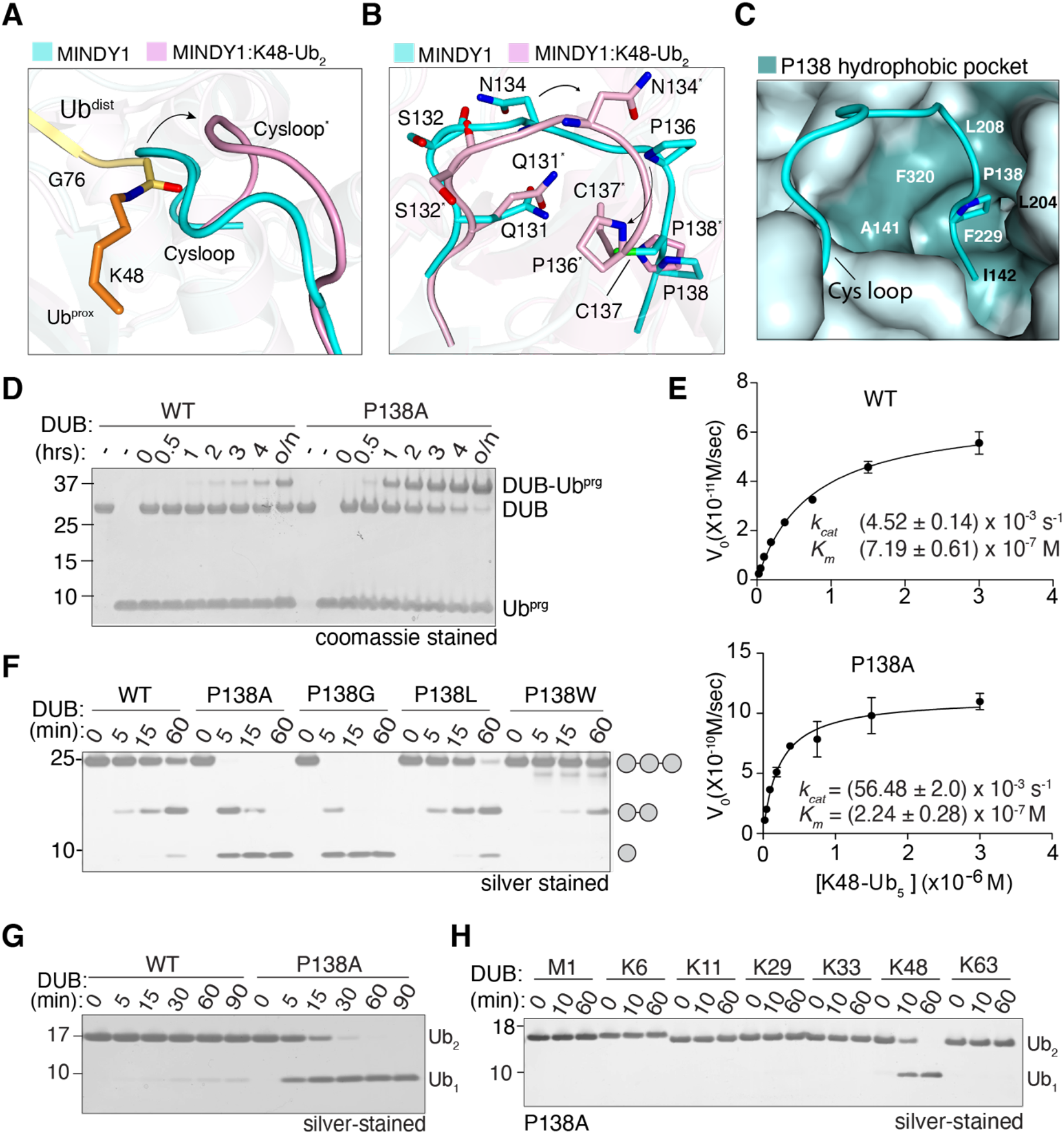
Cys loop mobility regulates DUB activity. **A**) Cartoon representation of the Cys loop in a superposition of MINDY1apo (cyan) and MINDY1:K48-Ub2 complex (pink). The isopeptide bond between K48 of Ub^prox^ (orange) and G76 of Ubdist (yellow) is shown in sticks. **B**) Close-up view of (**A**) showing amino acid side chain rearrangements (side view). **C**) Surface representation of the hydrophobic pocket in MINDY1apo that accommodates the Cys loop residue P138. **D**) Coomassie-stained gel comparing activity of MINDY1 WT and P138A to UbPrg in a time course. **E**) Steady-state kinetics of K48-linked pentaUb cleavage by MINDY1 WT and P138A mutant derived from reactions with varying concentrations of fluorescently-labelled Ub5. (n= 2; mean ± SD) **F**) Silver-stained gel comparing cleavage of K48-Ub3 by MINDY1 WT and indicated mutants. **G**) DUB assay comparing cleavage of K48-Ub2 by MINDY1 WT and P138A mutant. **H**) DUB assay monitoring cleavage of diUb of seven different linkage types by MINDY1 P138A.

The catalytic cysteine in both MINDY1 and MINDY2 are flanked by proline residues: MINDY1 (P136-C137-P138) and MINDY2 (P256-C266-P267). When we compare Cys loop conformations in the autoinhibited and active states of MINDY1, P138 in the Cys loop remains fixed while the rest of the loop moves (**Fig 2B**). P138 sits in a hydrophobic pocket formed by A141, I142, L204, L208, F229 and F320 to function as an anchor point for the Cys loop (**Fig 2C and S3C**). When in complex with diUb, the nature of the interactions of P138 within the hydrophobic pocket change with only L204, L208 and F320 mediating interactions, whereas F229 and I142 now no longer interact with P138 (**Fig S3D**). In contrast to the anchored P138, the other flanking proline, P136, shows significant movement upon complex formation. P136 does not form any intramolecular interactions in the apo state, however in the diUb-bound active state, P136 is part of a strong intramolecular interaction network consisting of hydrophobic interactions with Y114, L139, L140 and A205 (**Fig S3C,D**).

Based on these observations, we predict that P138 modulates the dynamics of the Cys-loop as it provides rigidity to the loop by anchoring itself into the hydrophobic pocket. Hence, mutation of P138 to alanine should dislodge the hydrophobic pocket interactions and result in a more dynamic, flexible loop. To test this possibility, we determined the crystal structures of MINDY1 P138A mutant on its own and in complex with K48-Ub2. In contrast to WT, electron density for the Cys loop in the P138A mutant has high b-factors, suggestive of a more mobile loop (**Fig S3 M,N**). To determine the effect of a more flexible Cys loop on DUB activity, we first assayed reactivity of MINDY1 towards propargylated ubiquitin (Ub^Prg^) that covalently modifies the catalytic Cys with the Ub occupying the S1 site. MINDY1 is not effectively modified by Ub^Prg^ with only about 50% modification after an overnight incubation. In contrast, the P138A mutant was readily modified by Ub^Prg^ with more than 50% conversion in 2 hours, thus supporting our notion that the P138A mutation makes the Cys loop more flexible and no longer sterically blocks ubiquitin binding (**Fig 2D**). Next, we determined enzyme kinetics using fluorescently labelled K48-linked pentaUb, which revealed that MINDY1 P138A mutant is a much more active enzyme with greater than 10-fold higher *k_cat_* and ~3-fold lower *K_m_* compared to WT (**Fig 2E**). As the Cys loop sterically interferes with polyUb binding, an increase in loop flexibility possibly might result in lower *K_m_*.

To test the role of the hydrophobic pocket in keeping P138 anchored, we mutated P138 to smaller residues (A or G), which would disrupt anchoring, or to bulky hydrophobic residues (L or W), which would lock the Cys loop in the inhibited state. A chain cleavage assay with these mutants shows that disrupted anchoring (P138A or P138G) increases chain cleavage whereas increased hydrophobicity of the side chain impairs DUB activity (**Fig 2F**). The consequence of a more flexible loop is also underscored by the ability of P138A to cleave diUb, which wildtype MINDY1 is unable to cleave (**Fig 2G**). Similarly, mutating the equivalent residue in MINDY2 (P267A) also leads to increased DUB activity (**Fig S3J, K**). Importantly, increasing Cys loop flexibility with the P138A mutation, despite making the enzyme more active, does not change its linkage-specificity for K48 chains as the mutant only cleaves K48 chains and none of the other linkage types (**Fig 2H**). Collectively, these results demonstrate that the Cys loop regulates MINDY1 catalytic activity but not linkage selectivity.

### Mechanism of autoinhibition

In comparison to P138, the MINDY1 P136A mutant shows marginally reduced activity compared to WT (data not shown). To analyse this further, we introduced the P136A mutation onto the MINDY1 P138A mutant. When activity was assayed using K48-Ub_2_ as substrate, the P136A mutant, like WT, is unable to cleave diUb whereas the P138A mutant is very active (**Fig 3A**). However, the double mutant P136A/P138A shows diminished activity compared to the P138A mutant suggesting catalytic roles for P136. One notable interaction of P136 is with the phenyl ring of Y114 (**Fig S3D**). In the autoinhibited state of MINDY1, a sulphur-centred hydrogen bonding of the catalytic C137 with the hydroxyl of Y114 rotates C137 away from the active site (**Fig 3B**). In the active conformation, this inhibitory Y114-C137 interaction is broken due to the Cys loop movement, and the side chain of Y114 moves to now hydrogen bond with S163 (**Fig 3C**). Interestingly, in the MINDY1:Ub^Prg^ complex, which represents the product-bound intermediate state, Y114 is hydrogen bonded to S163 prior to the transition back from the activated to the inhibited state after ubiquitin chain hydrolysis (**Fig S4A**). These observations strongly imply a linchpin role for Y114 in regulating the switch from an inhibited to active state.

**Figure 3.**
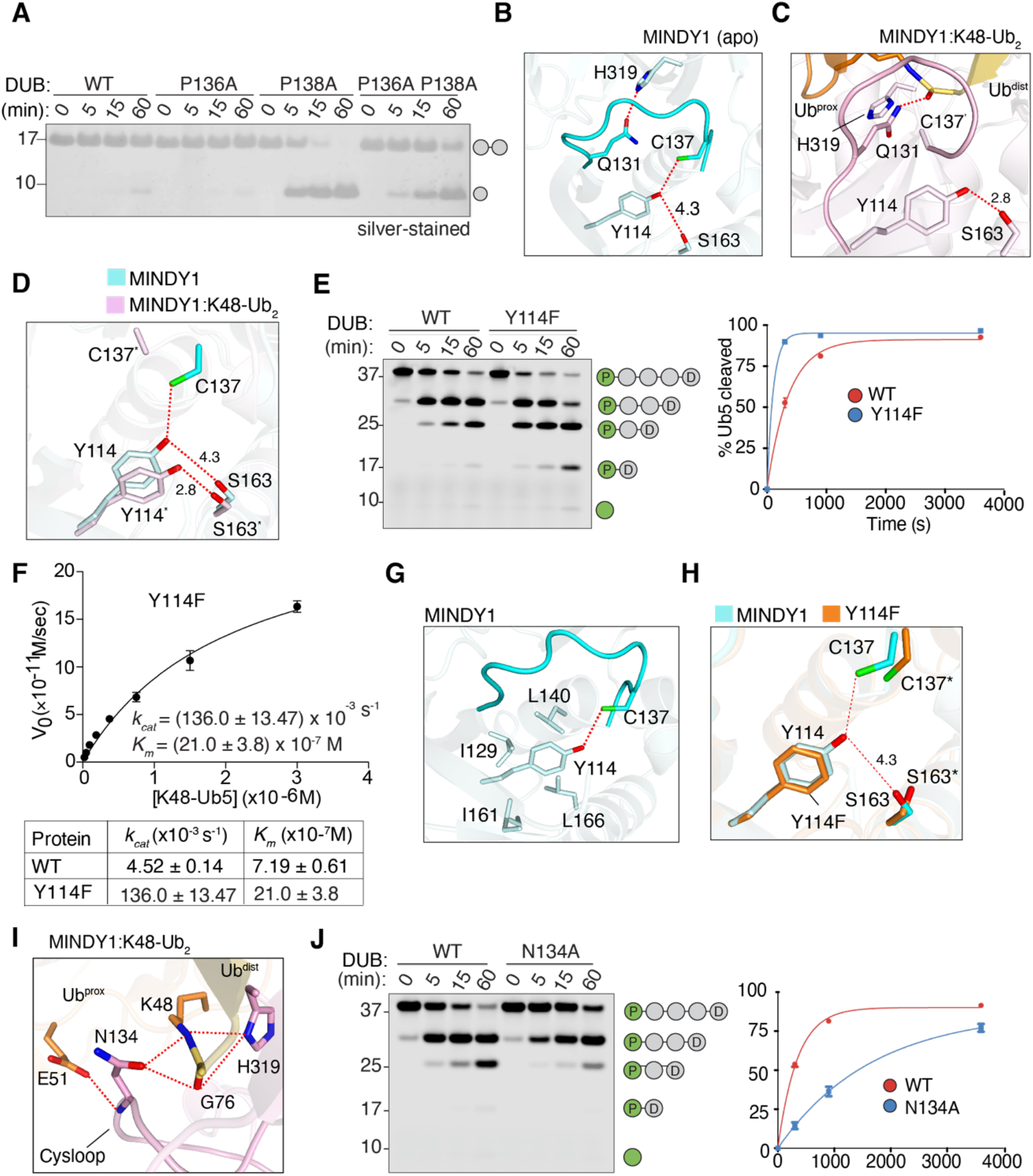
Autoinhibition and activation of MINDY1. **A**) DUB assay monitoring the cleavage of K48-Ub2 by MINDY1 and indicated mutants. **B**) Close-up view of catalytic residues and their interactions in MINDY1 (apo). C137 is out of plane with H139 and is hydrogen bonded with Y114 in MINDY1 (apo). Dotted lines in red indicate hydrogen bonds. **C**) Close-up view as in (**B**) for the MINDY1:K48-Ub2 complex. The catalytically productive state conformation leads to the formation of new sets of bonds as shown. The oxyanion hole residue Q131 which was in contact with catalytic H319 in (**B**) now forms interactions with the carbonyl of the incoming scissile bond. **D**) Lateral movement of Y114 and its interactions in MINDY1 (apo) and MINDY1:K48-Ub2 complex. **E**) DUB assays comparing cleavage of fluorescently-labelled pentaUb by MINDY1 and Y114F mutant. The percentage hydrolysis of pentaUb is plotted against time (right). **F**) Steady-state kinetics of K48-linked pentaUb cleavage by MINDY1 Y114F. (n= 2; mean ± SD) **G**) Close-up view of Y114 (phenyl ring) interactions with hydrophobic residues on adjoining secondary structure elements in MINDY1 (apo). **H**) A close-up image of active site of apo MINDY1 Y114F mutant compared to WT. Hydrogen bonding of C137 to Y114 is broken in the mutant. **I**) Interactions of Cys loop residue N134 in stabilizing the isopeptide bond for catalysis. **J**) Cleavage of pentaUb chains fluorescently-labelled on Ubprox by MINDY1 WT and N134A mutant. The panel on the right show the quantification of the DUB assay.

To test the importance of Y114 in keeping MINDY1 in an inhibited state, we introduced a Y114F mutation to disrupt the inhibitory interaction with the catalytic Cys. Indeed, loss of this single hydrogen bond results in increased DUB activity with ~30-fold increase in *k_cat_* compared to WT (**Fig 3E-H**). A conserved mechanism operates in MINDY2 as mutation of the analogous residue Y243 to F also leads to increased DUB activity (**Fig S4B-C**). Y114 is surrounded by hydrophobic residues which interact and stabilize the position of the phenyl ring (**Fig 3G**). Indeed, in the crystal structure of the MINDY1 Y114F mutant, the phenyl ring of F114 occupies the same position as Y114 of inactive MINDY1 (**Fig 3H**). On mutation of Y114 to alanine, MINDY1 is less active compared to Y114F mutant (**Fig S4E**). Of note, expression and stability of MINDY1 is affected by the Y114A mutation suggesting an additional structural role for Y114 in stabilizing local structure especially around the Cys loop. Interestingly, the equivalent residue of S163 of MINDY1 is an alanine (A292) in MINDY2 which is evolutionarily conserved (**Fig S4A**). Hence, the lateral movement of the side chain of Y114 observed during release of inhibition in MINDY1 does not occur for Y243 in MINDY2 (**Fig S4B**). However, there is a displacement of the catalytic C266 when MINDY2 is in complex with substrate, which breaks the interaction with Y243. Overall, these results highlight the importance of a non-catalytic tyrosine residue in regulating the Cys loop dynamics and thus regulating the activity of MINDY1 and MINDY2.

In addition to P138 and Y114, the inhibitory conformation of the Cys loop is further maintained by the intramolecular interaction between N134 and S132 in MINDY1. (**Fig S3B**). When in complex with K48-Ub_2_, this interaction is broken as S132 now contacts Q49 and R42 of Ub^prox^ and N134 interacts with K48, G76 and E51 of Ub^prox^ (**Fig 3I**). These interactions of the side chain of N134 with substrate are significant for deubiquitylation as mutating N134 to alanine impairs the catalytic activity of MINDY1 (**Fig 3J**). This suggests a model of substrate-assisted catalysis where interaction of S132 and N134 of MINDY1 with residues in ubiquitin enable the transition of MINDY1 from an inhibited to an active enzyme.

In the autoinhibited state of MINDY1, the oxyanion hole forming Q131 is hydrogen bonded to the catalytic H319. In the active state observed in the MINDY1: K48-Ub_2_ complex, N134 and the catalytic H319 both form hydrogen bonds with K48. (**Fig 3B, C**). In addition to the interactions with Ub^prox^ that stabilize the catalytic site in a productive conformation, Ub^prox^ also stabilises the binding of Ub^dist^ onto the S1 site (**Fig S2C**). Hence, in a substrate-activated mechanism, the binding of K48-linked polyubiquitin stabilizes the productive conformation of the catalytic site in MINDY1 and MINDY2. Indeed, in the absence of Ub^prox^, the active site in the MINDY1:Ub^Prg^ structure, shows both H319 and Q131 to exist in two alternate conformations. In one conformation, H319 is hydrogen bonded to Q131 similar to the inhibited state, and in another conformation, Q131 is flipped out and does not interact with any residue (**Fig S4H**). In contrast, in MINDY2^apo^, the catalytic H448 is flipped out, and Q260 is not able to form a hydrogen bond (**Fig S4J**). However, the binding of Ub^prox^ brings H448 closer to the catalytic C266 to form a productive active site (**Fig S4K**). In summary, the Cys loop dynamics and the interactions with K48-linked chains modulate the transition of MINDY1 and MINDY2 from an inhibited to active state.

### Non-canonical catalytic mechanism

In most thiol proteases, a third catalytic residue, usually an Asp or Asn, serves to correctly position the catalytic His (Clague *et al*, 2019). In all determined structures of MINDY1 and MINDY2, the identity of this third catalytic residue is unclear. In the MINDY1:K48-Ub_2_ structure, S321 is 3.6 Å away from H319 and could function as a catalytic residue as described for USP30 (**Fig 4A**) (Gersch *et al*, 2017; Sato *et al*, 2017). However, to our surprise, mutating S321 to Ala did not abolish activity but instead resulted in a modest increase in activity (**Fig 4D**). A search for other potential residues only revealed a distant T335 situated approximately 6 Å away (**Fig 4A**).

**Figure 4.**
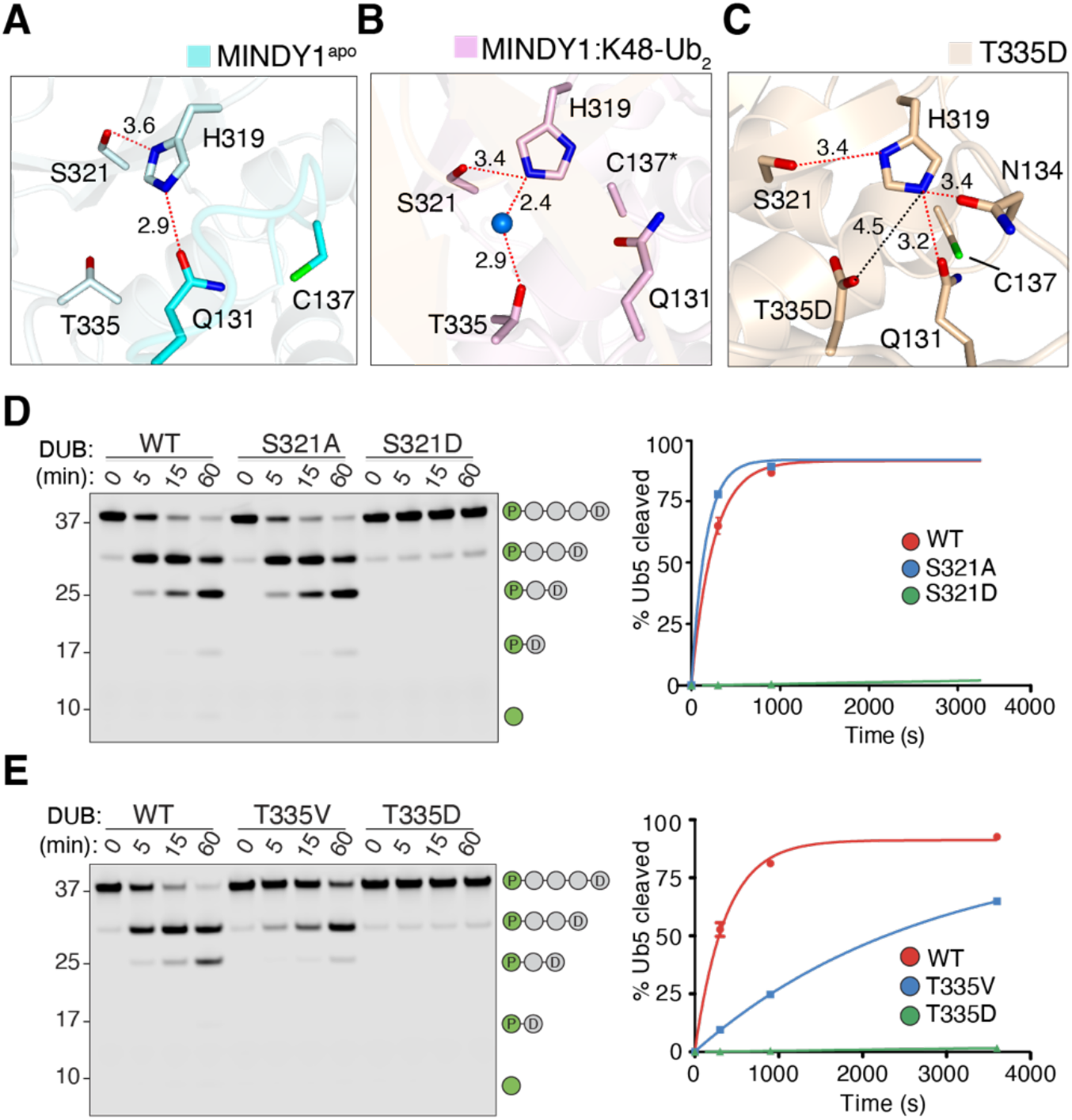
MINDY1 uses a non-canonical catalytic mechanism. **A**)-**C**) Close up view of the catalytic site in MINDY1 apo (**A**), MINDY1:K48-Ub2 complex (**B**) and in MINDY1 T335D mutant (**C**). Dotted red lines indicate hydrogen bonds, dotted black line ionic bond and blue sphere indicates water molecule. **D**) DUB assay comparing the cleavage of fluorescently labelled K48-Ub5 by MINDY1 WT, S321A and S321D mutants. The percentage of pentaUb hydrolysed is plotted against time (right). **E**) DUB assay as in (**D**) comparing the chain cleavage by MINDY1 WT, T335V and T335D mutants. The percentage of pentaUb hydrolysed is plotted against time (right).

Despite this distance, a water molecule coordinates T335 with H319 via hydrogen bonding and may serve to orient H319 for catalysis, thereby adopting the function of the third catalytic residue (**Fig 4B**). To test if such an unusual catalytic architecture is possible, we introduced a T335V mutation to disrupt this water bridge, which resulted in substantial loss of activity (**Fig 4E**). Hence, T335 competes with S321 to correctly position H319 for catalysis, thus adding another layer of regulation. S321 thus functions as an inhibitory residue and this explains why the S321A mutation enhances activity. Further supporting this model is the mutation of S321 to aspartate which completely abolishes the catalytic activity, as a strong ionic bond between S321D and the catalytic histidine (H319) likely blocks the catalytic function of H319 thus rendering the DUB inactive (**Fig 4D**).

T335 therefore serves a catalytic function to correctly position H319 for catalysis. We therefore wondered if mutating T335 to a negatively charged residue would enhance the activity of MINDY1. To our surprise, the mutation of T335 to aspartate also abolishes MINDY1 activity (**Fig 4E**). To understand the reason underlying this loss of activity, we determined the crystal structure of MINDY1^T335D^ (PDB ID: 6YTR) which revealed the formation of an ionic interaction between T335D and the catalytic H319. H319 is locked in this conformation which is further strengthened by the network of hydrogen bonds with Q131 and N134 resulting in the catalytic Cys being rotated away from H319 (**Fig 4C, S5A**). Hence mutation of T335 to aspartate also forces the enzyme into an unproductive catalytic state, thus highlighting the requirement of a water molecule bridged hydrogen bonding interaction with T335 to correctly position H319 for an active enzyme. MINDY2 also uses a similar non-canonical catalytic architecture where T464 is the third catalytic residue and S450 serves as a competitive element (**Fig S5B-D**).

### Why does MINDY1 preferentially cleave long Ub chains?

The minimal catalytic domains of both MINDY1 and MINDY2 show higher activity at cleaving longer chains as revealed by DUB assays with fluorescently-labelled polyubiquitin chains of increasing length (**Fig 5A, B**). We hypothesize that the preference of these DUBs for cleaving longer chains is due to the presence of additional Ub binding sites within the catalytic domain. Indeed, unexplained electron density adjacent to α3-α5 (near the S1 site) in the MINDY1: K48-Ub_2_ crystal structure revealed it to be that of an additional ubiquitin moiety. However, this lies at the symmetry axis and only a partial ubiquitin molecule could be built (**Fig S6D**). Compared to MINDY1, the crystal structure of MINDY2:K48-Ub_2_ contains two complexes in the ASU. Unlike the MINDY1: K48-Ub_2_ complex, there is a fully traceable extra ubiquitin bound at this equivalent position in the MINDY2 complex (**Fig S6E**). Moreover, in one of the complexes in the ASU, there is an additional fourth ubiquitin bound at a helix-loop-helix region (α2-α3) of MINDY2 (**Fig S6F**). The presence of these extra ubiquitin densities in our structures may be due to monoubiquitin contamination in the sample used for crystallisation of the complex. Nevertheless, they suggest the presence of additional ubiquitin binding sites within the catalytic domain. If MINDY1 and MINDY2 do indeed have additional ubiquitin binding sites, we predict that they would have higher affinity for longer K48 chains compared to K48-Ub_2_. Hence, we performed isothermal calorimetry (ITC) measurements where catalytically dead MINDY1 C137A was titrated into K48-diUb, triUb, tetraUb or pentaUb (**Fig 5D**). To our surprise, there was no measurable binding for diUb by ITC where the chain would occupy the S1 and S1’ sites. In contrast, we observe an increase in binding affinity with increasing chain length. MINDY1 binds to triUb with an affinity of ~6μM ± 3μM which increases to ~1μM ± 0.2μM for tetraUb and a further increase to ~250nM for pentaUb. Further, the binding affinity for pentaUb determined by ITC (K_d_ ~ 250 nM) is consistent with the *K_m_* (*K_m_* =700 nM). These results indicate that MINDY1 has at least five distinct Ub binding sites within its catalytic domain and the stoichiometry of 1:1 in the ITC measurements suggests that one pentaUb chain is bound by a single molecule of MINDY1 (**Fig 5C**).

**Figure 5.**
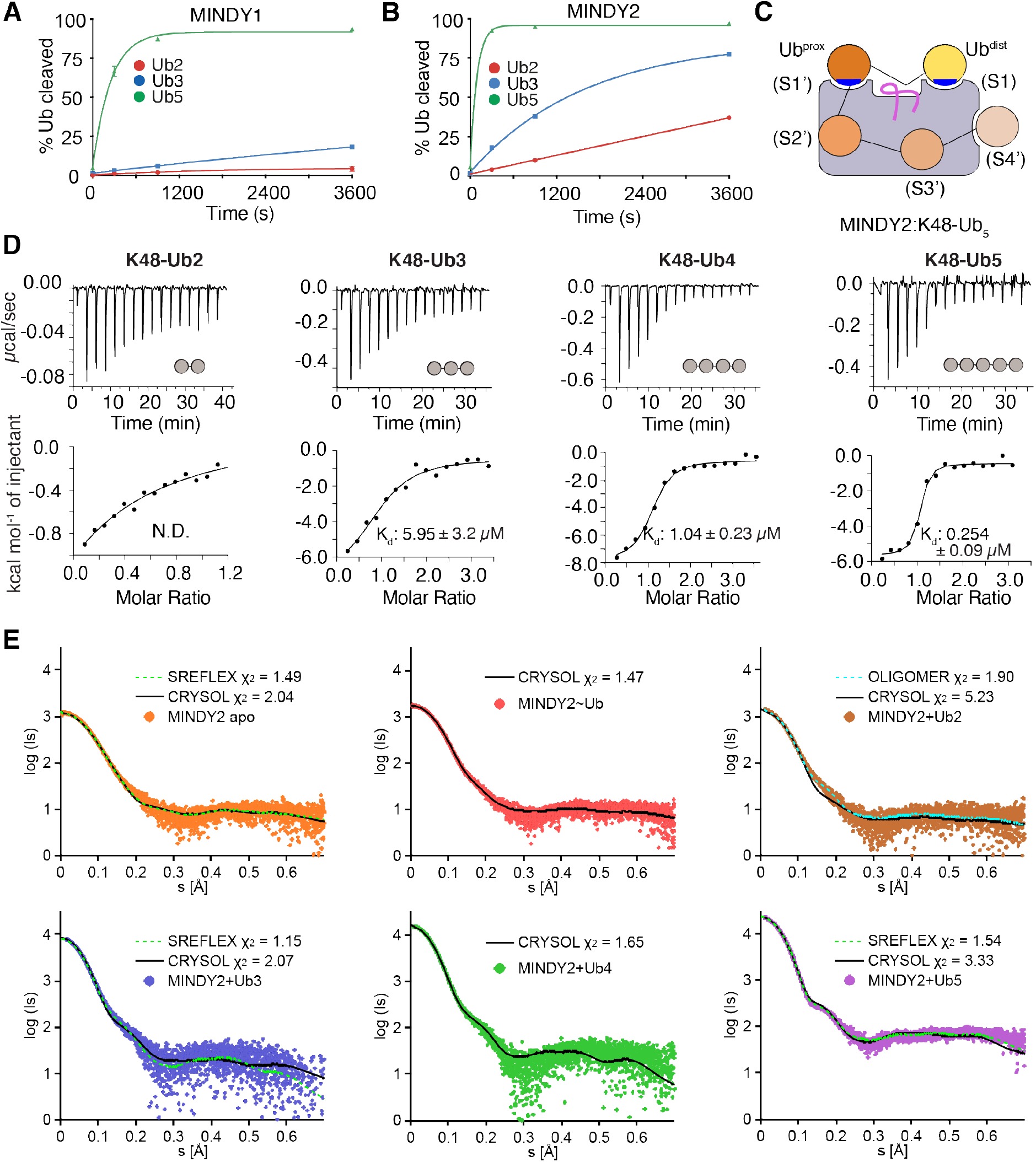
Five distinct Ub binding sites on the catalytic domain of MINDY1 and MINDY2. **A-B)** MINDY1 and MINDY2 prefer to cleave longer chains. The percentage of fluorescently-labelled K48-linked Ub2, Ub3 and Ub5 cleaved over time by MINDY1 and MINDY2 is shown. The data shown for K48-Ub5 cleavage by MINDY1 was plotted using the data from Fig 4D and 4E. (n = 2; mean ± SD). **C**) Model depicting ubiquitin chain bound at the S1, S1’ sites and the putative S2’, S3’ and S4’ binding sites. **D**) Isothermal titration calorimetry (ITC) data measuring binding of MINDY1cat with K48-linked polyubiquitin chains of the indicated lengths. **E**) SAXS curves of MINDY2, apo molecule and in complex with monoUb, K48-linked Ub2, Ub3, Ub4 and Ub5, respectively, and their fits computed from atomic models by CRYSOL. For Mindy2-Ub3 and Mindy2-Ub5, NMA with SREFLEX was used for the refinement of the expected atomic models. For Mindy2-Ub2, OLIGOMER was used on atomic models with Ub2 occupying positions S1 and S1’ or S1 and S3’ to quantify their mixture.

Despite several attempts, we were unable to obtain crystals of MINDY1 or MINDY2 in complex with K48-linked tetraUb or pentaUb. To validate the presence of additional ubiquitin binding sites on MINDY1 and MINDY2, we therefore performed small-angle X-ray scattering (SAXS) on MINDY2 complexes. SAXS data were collected for MINDY2 on its own, covalently linked to Ub^Prg^ and in complex with K48-Ub_2_, Ub_3_, Ub_4_ and Ub_5_ (**Table S4**). MINDY2 apo and its complexes were chosen for SAXS analyses because of the ordered ubiquitin density seen at the putative fourth and fifth binding site in the crystal structure of the MINDY2 (**Fig S6F**). These structures provided the initial models for MINDY2: K48-Ub_3_, MINDY2: K48-Ub_4_ and MINDY2:K48-Ub_5_ for the SAXS study (**Fig 5C, S6A**).

The SAXS curves were first computed from all the working models mentioned above by using CRYSOL (Svergun et al., 1995) and compared with the experimental data. For both MINDY2 apo and MINDY2:Ub^Prg^ complex, computed data from the models yielded good agreement with the experiment confirming the two structures in solution (**Fig 5E and Table S4**). For complexes with ubiquitin chains, some deviations were observed (**Fig 5E, S6B-C**), and the expected models were further refined to fit the SAXS data. Thus, for MINDY2:K48-Ub_3_ normal mode analysis (NMA) was performed using SREFLEX (Panjkovich & Svergun, 2016) yielding a conformation close to the predicted model (RMSD of 5.01 Å). The SAXS data of the MINDY2:K48-Ub_2_ complex was analysed with the program OLIGOMER (Konarev *et al*, 2003) to check for the presence of complexes with distinguishable Ub_2_ binding positions. The best fit was obtained for a mixture of two different binding modes: productive (~15%), where ubiquitins are bound in the S1 and S1’ sites and unproductive (~85 %), where ubiquitins are bound in the S1 and S3’ sites (**Fig 5E, S6B**). The higher prevalence of ubiquitin binding in the unproductive mode possibly explains the inefficient cleavage of K48-Ub_2_ by MINDY1 and MINDY2. Interestingly, both MINDY1:K48-Ub_2_ and MINDY2:K48-Ub_2_ were crystallised in the productive state due to crystal packing where the unproductive binding sites revealed by SAXS are blocked by crystal contacts made with adjacent DUB molecules in the crystal lattice. Moreover, in both MINDY1:K48-Ub_2_ and MINDY2:K48-Ub_2_ structures, MINDY1 and MINDY2 form dimers via a disulphide bond (C358 in MINDY1 and C429 in MINDY2) (**Fig S6G**). SAXS measurements for the MINDY2:K48-Ub_3_ complex agree reasonably well with the proposed model and thus give us an idea of the putative S2’ site (**Fig S6C**). The MINDY2:K48-Ub_4_ SAXS measurements show that the ubiquitin molecules occupy the S1, S2, S1’ and S2’ sites (**Fig S6C**). Importantly, for MINDY2:K48-Ub_5_ complex a good agreement with the experiment is obtained by a mild NMA refinement (RMSD 2.12 Å to the proposed model revealing the S3’ and S4’ sites, **Fig 5E**). Put together, the SAXS data reveals the possible location of the S2’, S3’ and S4’ binding sites on MINDY2^cat^ (**Fig 5C**). To evaluate the importance of the five distinct Ub binding sites to the activity of MINDY1, we mutated residues that based on the SAXS data would form the S4’ site. This revealed that mutating the S4’ site results in impaired cleavage of K48-linked pentaUb (**Fig S6H**). In summary, the enzyme assays, ITC and SAXS studies support our hypothesis that five distinct ubiquitin binding sites are present within the minimal catalytic domain, thus explaining the preference of both MINDY1 and MINDY2 for cleaving pentaUb chains.

### Ubiquitin chain length determines a switch between exo and endo cleavage activities

We had previously demonstrated that MINDY1 is an exo-DUB that cleaves pentaUb from the distal end of the chain (Abdul Rehman *et al*, 2016). One possible mechanism to achieve such directionality of cleavage would be that MINDY1 recognizes the distal end of the chain. However, close examination of the MINDY1:K48-Ub_2_ and MINDY2:K48-Ub_2_ structures revealed that neither K48 and surrounding residues on Ub^dist^ nor G76 of Ub^prox^ mediate intermolecular contacts with MINDY1/2 (**Fig S7A, B**). The solvent exposed K48 on Ub^dist^ implies that this Ub does not necessarily have to be the extreme distal moiety in a chain. MINDY1/2 could therefore in principle possess endo-activity and cleave within a ubiquitin chain. Since all our previous assays were performed with pentaUb and our identification of five ubiquitin binding sites within the catalytic domain, we carefully analysed how MINDY1/2 would cleave chains containing more than five Ub moieties.

On incubation with polyUb chains, an exo-DUB will cleave one ubiquitin moiety from the end of the chain. Thus, a characteristic of an exo-DUB is the appearance of monoubiquitin as a predominant product at the earliest time points of a chain cleavage assay. Only at later stages, as the chain is gradually trimmed, do shorter chains such as diUb appear. In contrast, a DUB with endo-activity can cleave anywhere within a polyubiquitin chain, resulting in the formation of chains of all intermediate lengths as products. When long K48-linked chains containing more than five Ub moieties were incubated with MINDY1/2, the long chains collapsed to predominantly a mixture of chains containing up to four Ub molecules (**Fig 6A, B**). Compared to the long chains, tetraUb is not a preferred substrate as it rapidly accumulates followed by gradual cleavage only at later time points. We further confirmed this conditional endo-DUB activity using defined hexaUb as a substrate which results in both Ub5 and Ub4 being generated as the first products (**Fig 6C, D**). These results suggest that with chains longer than pentaUb, MINDY1/2 can work as an endoDUB to cleave within the chain, whilst with shorter chains, MINDY1/2 works as an exo-DUB to cleave the distal Ub. These assays reinforce the preference for the occupation of all five binding sites and offer a potential explanation for the switch between endo- and exo-cleavage. The most favourable binding mode that results in efficient cleavage is that all Ub-binding sites on the DUB are occupied. Therefore, MINDY1/2 can efficiently bind a K48-ubiquitin chain of length n (with n≥5) in n-4 possible ways within the chain thus leading to a rapid collapse of longer chains to tetraUb. As the chains get shorter (n<5), all five binding sites can no longer be occupied, thus leading to a decrease in cleavage efficiency. TetraUb for instance does not occupy all binding sites on the DUB and consequently cleavage is inefficient resulting in the observed accumulation of Ub_4_. With n=5, the minimum chain length that can satisfy the binding requirements, an exo-form of cleavage is forced with one Ub monomer being trimmed from the chain with each cleavage event. Hence, in addition to being specific for K48-linked polyUb, MINDY1 and MINDY2 also sense ubiquitin chain length to position and cleave long K48-linked chains down to tetraUb.

**Figure 6.**
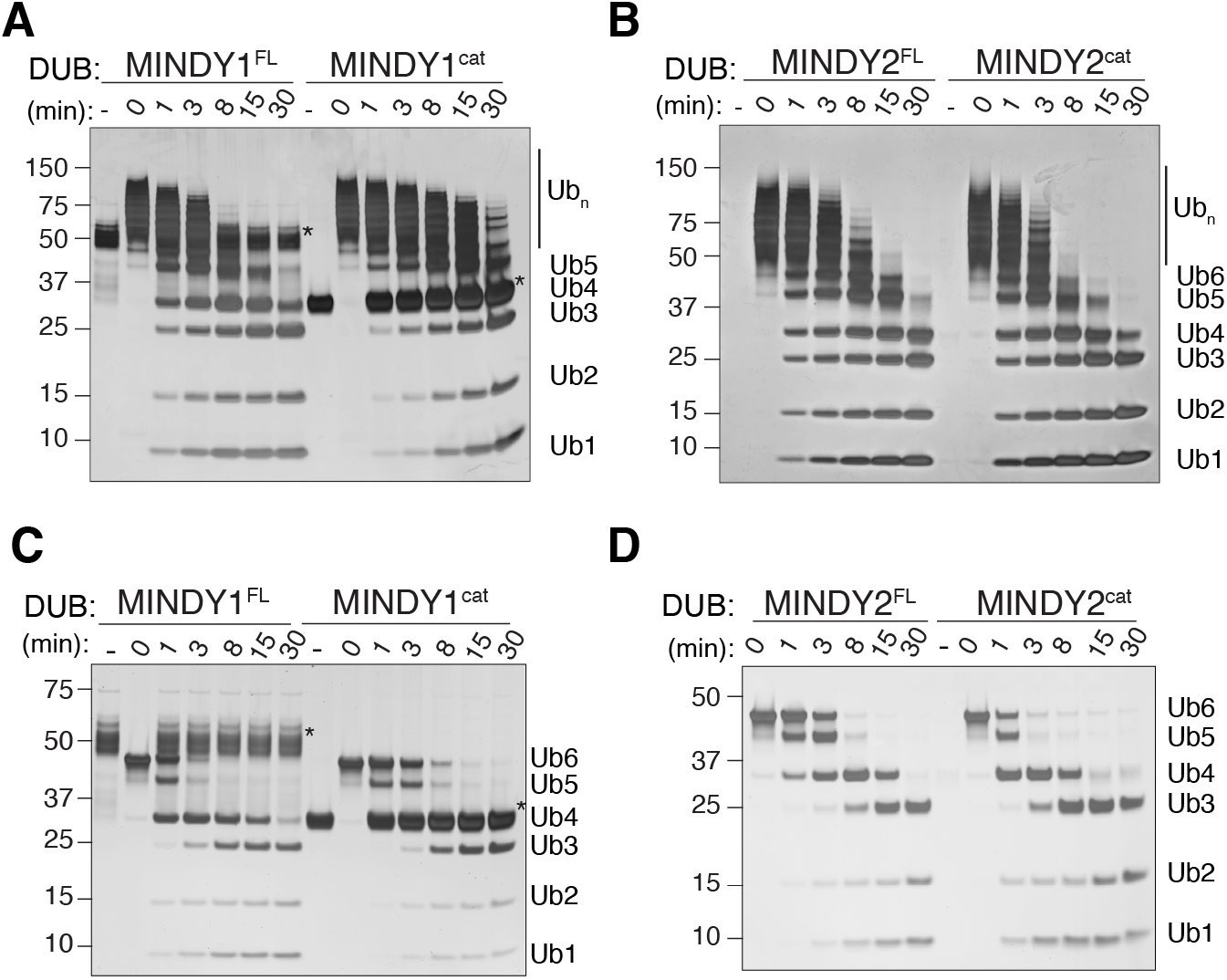
Ubiquitin chain length determines a switch between exo and endo cleavage activities. **A**) Silver-stained gels of DUB assays monitoring cleavage of long K48-linked polyUb chains containing more than 6 Ub moieties by full length and the minimal catalytic domain of MINDY1. **B**) DUB assay as in A) comparing cleavage by full length and catalytic domain of MINDY2. **C**) DUB assay as in (A) of a time course of K48-linked Ub6 cleavage by MINDY1^FL^ and MINDY1^cat^. **D**) DUB assay as in (B) of a time course of K48-linked Ub6 cleavage by MINDY2^FL^ and MINDY2^cat^. Control reaction containing only the DUB is shown in the lane (-) and the lane corresponding to 0 min has only polyUb chain before addition of DUB. Asterisk indicates MINDY1FL and MINDY1cat protein.

## DISCUSSION

The three crystal structures of MINDY1 reveal distinct states in the catalytic cycle of MINDY1, namely: autoinhibited (Apo), substrate-bound active state (MINDY1:K48-Ub_2_) and the product-bound intermediate state (MINDY1:Ub^Prg^) (**Fig 7, S7C**). In the apo state, the Cys loop mediates autoinhibition and sterically interferes with ubiquitin binding. While large scale conformational changes are not observed upon enzyme:substrate complex formation, ubiquitin binding at the S1’ site releases the autoinhibition mediated by the Cys loop to enable ubiquitin binding at the S1 site. Hence, in a model of substrate-driven activation, ubiquitin bound at the S1’ site interacts with the Cys loop to drive the transition of MINDY1 and MINDY2 from inhibited to active enzymes. The remodelling of the Cys loop also brings the catalytic residues into a productive conformation to enable catalysis.

**Figure 7.**
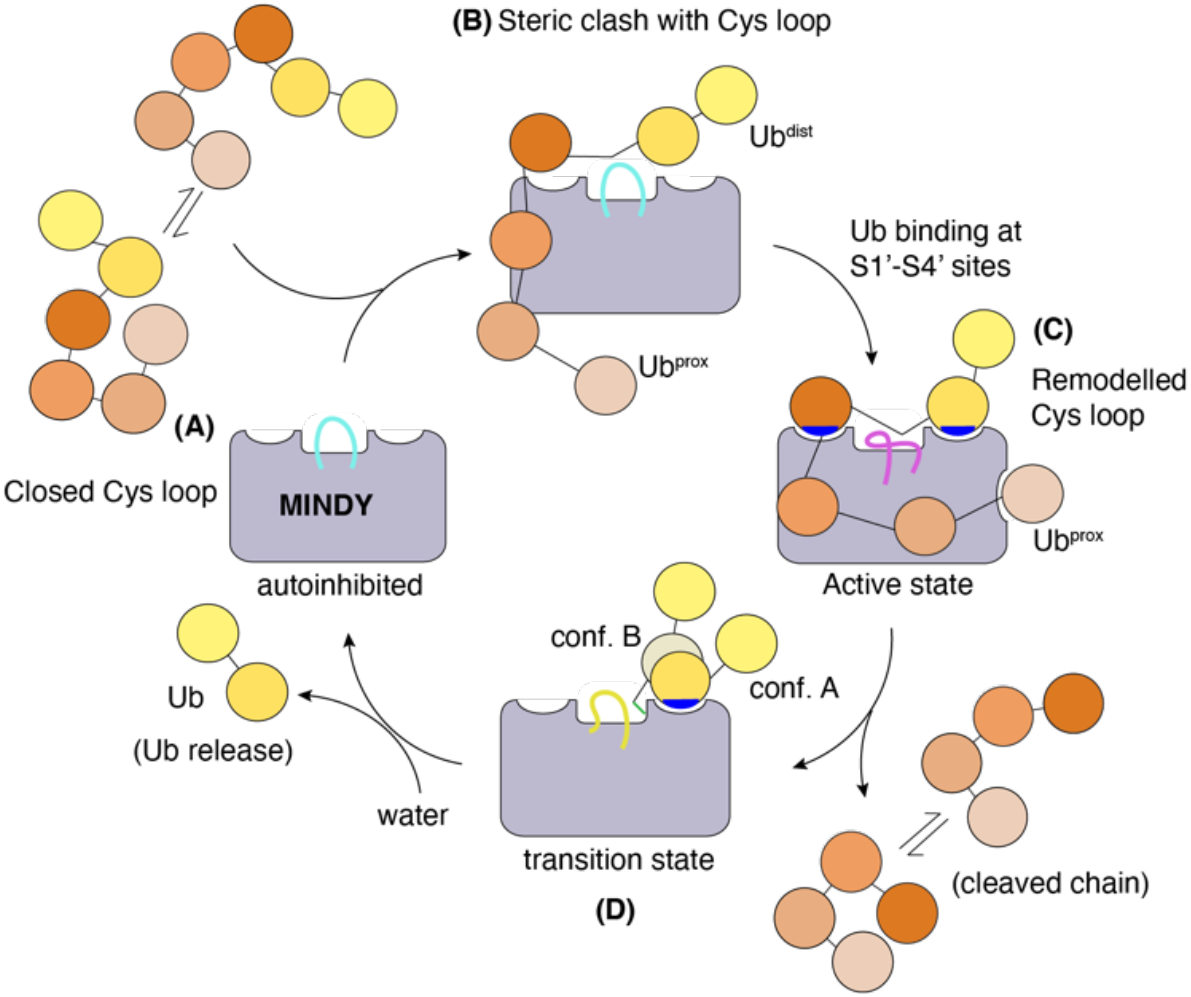
Model summarising the catalytic mechanism of K48 linked polyubiquitin. MINDY1 and MINDY2 exist in an autoinhibited conformation where the Cys loop is in a closed conformation which sterically interferes with ubiquitin binding and also contributes to keeping the catalytic site inhibited (**A**). Ubiquitin chains exist in a dynamic equilibrium between closed and open conformations. Cys loop remodelling requires a ubiquitin chain to bind and occupy all five sites on the catalytic domain. In a substrate-driven mechanism, ubiquitin interactions release inhibition and activate the DUB resulting in chain cleavage and release of the ubiquitin chain (**B**-**C**). In the product intermediate transitional tetrahedral state, the ubiquitin occupies the S1 site. As this is not a strong binding interface this ubiquitin exists in two different conformers. Attack by a water molecule releases the ubiquitin and returns the DUB to an inhibited conformation.

In MINDY1 and MINDY2, the correct positioning of the scissile bond requires Ub binding at both the S1 and S1’ sites. In several DUBs, polyUb recognition and cleavage depends on extensive Ub interactions at the S1 site (Leznicki & Kulathu, 2017). When binding at the S1 site is not very strong, DUBs then rely on additional interactions at the S1’ site. Indeed, this is a hallmark of DUBs such as CYLD and OTU family DUBs where ubiquitin interactions at the S1’ site places the proximal Ub in a position that orients a specific lysine into the catalytic site, thus conferring linkage specificity (Licchesi *et al*, 2012; Mevissen *et al*, 2016; Sato *et al*, 2015). A similar mechanism drives linkage-selectivity in MINDY1 and MINDY2, where proximal Ub interactions position it such a way that only K48-linked polyubiquitin can access the catalytic groove. However, the simultaneous engagement of polyUb at the S1 and S1’ sites we think also requires binding of ubiquitins at the S2’-S4’ sites since, in their absence, the SAXS data indicates that a significant proportion of diUb binds the DUB in a non-productive conformation at the S1 and S3’ sites. MINDY1 has weak affinity for K48-linked diUb and the steric hindrance imposed by the Cys loop prevents diUb binding. The P138A mutation which increases the mobility of the Cys loop not only enables transition to the active conformation but also no longer interferes with ubiquitin binding as evidenced by lower *Km* for pentaUb and binding to diUb. Further, only the P138A mutant is able to cleave diUb which MINDY1 WT cannot cleave, which leads us to suggest that efficient remodelling of the Cys loop occurs when pentaUb is bound by the DUB. The high binding affinity that pentaUb has for the catalytic domain and the presence of 5 distinct Ub binding sites within the catalytic domain makes MINDY1/2 unique amongst DUBs.

Free K48-linked chains predominantly exist in a closed conformation as hydrophobic interactions between I44 patches of the ubiquitin moieties form a tight binding interface and this compact conformation cannot be recognized by DUBs (Cook *et al*, 1992; Eddins *et al*, 2007; Trempe *et al*, 2010). In addition, “open” forms of K48 chains have also been observed (Hirano *et al*, 2011; Lai *et al*, 2012; Lu *et al*, 2020). Polyubiquitin chains including K48-linked chains are dynamic in solution and exist in dynamic equilibrium between closed and open conformations (Hirano *et al*, 2011; Ye *et al*, 2012). Our structure of MINDY1 and MINDY2 in complex with K48-linked diUb reveals that the K48 chains adopt an extended conformation. Moreover, both the proximal and distal ubiquitin interact with MINDY1/2 using their I44 patches, a binding mode that results in the K48-linked chain adopting a unique conformation not previously observed. Since MINDY^cat^ binds to longer chains, it is not clear if these long chains containing more than five Ub moieties more readily adopt an open conformation compared to diUb, thus favouring recognition and cleavage. Alternatively, an initial binding event involving the I44 patch of one ubiquitin could trigger conformational rearrangement in the remainder of the chain.

We previously reported that MINDY1 is an exo-DUB that sequentially cleaves Ub one at a time from the distal end (Abdul Rehman *et al*, 2016). This conclusion was drawn from experiments using pentaUb as substrate. Using longer Ub chains as substrate, we here clearly demonstrate that MINDY1 and MINDY2 exhibit endo- or exo-DUB cleavage that is dependent on the chain length. With long polyUb chains, MINDY1/2 recognize and bind blocks of pentaUb to cleave as an endo-DUB, thus releasing free Ub chains as product. With pentaUb, the extreme distal ubiquitin in the chain is positioned at the S1 site of the DUB, thus forcing an exo-mode of cleavage in which one moiety is trimmed at a time. The lack of any mechanism by which the distal end of the Ub chain is specifically recognized further reinforces MINDY1 and MINDY2 as endo-DUBs that recognize and cleave long ubiquitin chains, such that substrates are left with tetraUb chains. An emerging theme in ubiquitylation is a role for the length of the polyUb chain in determining the consequence. For instance, the protease DDI2 will cleave the transcription factor NRF1 only when it is modified with long polyUb chains (Dirac-Svejstrup *et al*, 2020). Similarly, the unfoldase Cdc48/p97-Ufd1-Npl4 has several ubiquitin binding sites and efficiently unfolds Mcm7 only when it is modified with K48 chains containing atleast five ubiquitins (Deegan *et al*, 2020; Twomey *et al*, 2019).

Our studies of MINDY1 and MINDY2 reveal two very similar enzymes, with similar regulatory mechanisms and substrate specificities. This provokes the question as to why evolution would preserve and maintain two relatively similar DUBs with similar properties. MINDY1 and MINDY2 are modular DUBs that in addition to their catalytic domain possess tandem MIU domains (tMIU), which may aid in substrate recruitment and cleaving longer polyubiquitin chains (Abdul Rehman *et al*, 2016). The tMIU of MINDY1 is highly selective at binding to K48-linked polyubiquitin chains, with MIU2 of the tandem motif being the main determinant of this specificity (Kristariyanto *et al*, 2017). Interestingly, the tMIU of MINDY2 is non-specific and binds to polyubiquitin chains of different linkage types. This raises the possibility that MINDY2 cleaves polyubiquitin containing mixed or branched ubiquitin linkages, where the tMIU binds to the non-K48-linked part of the chain and the DUB cleaves the K48 linkages, suggestive of a distinct cellular function from MINDY1.

MINDY1 and MINDY2 exist in an autoinhibited conformation characterized by misaligned catalytic residues and a Cys loop that impedes ubiquitin binding. Crystal structures of many DUBs reveal that their catalytic residues are often in unproductive conformations in the absence of substrate and conformational rearrangements are triggered by ubiquitin binding that lead to realignment of the catalytic residues into a productive conformation (Hu *et al*, 2002; Mevissen *et al*, 2016; Keusekotten *et al*, 2013; Sato *et al*, 2015; Boudreaux *et al*, 2010). In MINDY1/2 ubiquitin binding results in several rearrangements leading to the formation of a functional active site. Most thiol DUBs feature a canonical catalytic triad composed of Cys, His and Asp/Asn (Ronau *et al*, 2016). The Asp/Asn plays a secondary role by properly orienting the His in the catalytic triad. DUBs such as USP16, USP30 and USP45 have a serine in place of the Asp/Asn making them distinct from other DUBs (Gersch *et al*, 2017; Sato *et al*, 2017). MINDY1 features an atypical catalytic triad where a Thr residue orients the His via a water bridge. Intriguingly, a local non-catalytic Ser residue plays an inhibitory role by competing with the catalytic Thr for interaction with the catalytic His and improperly orienting it. This is reminiscent of OTULIN where inhibitory interactions mediated by an Asp with the catalytic His inhibit the DUB and are relieved upon substrate binding (Keusekotten *et al*, 2013). In MINDY1 and MINDY2, an additional layer of regulation is imparted by a sulphur-centred hydrogen bond between a Tyr and the catalytic Cys which further reinforces autoinhibition.

The mobility of the inhibitory Cys loop in MINDY1 is restricted by the side chain of P138 which is anchored into a hydrophobic pocket. Disrupting this anchoring by mutating P138 to a smaller residue leads to increase activity whereas a bulkier residue inhibits activity. Miy2 (Ypl191C), the yeast ortholog of MINDY2, has an alanine at this position (A29) and may explain why it is a much more active enzyme compared to MINDY1 (Abdul Rehman *et al*, 2016). The other flanking proline in the Cys loop P136 has a catalytic role and its mutation impairs DUB activity. Intriguingly, in Miy1 (Ygl082W) which does not cleave ubiquitin chains despite having all catalytic residues, the equivalent residue of P136 is a glutamate (E27) (Abdul Rehman *et al*, 2016). In summary, our work reveals that MINDY1 and MINDY2 are unique DUBs that sense both ubiquitin chain length and linkage type. The remarkable specificity that MINDY1 and MINDY2 possess at cleaving K48-linked chains to trim long polyubiquitin chains may help reveal the cellular functions of these evolutionarily conserved DUBs. That MINDY1 and MINDY2 have evolved so many layers of regulation and activation steps suggests key regulatory functions for these DUBs.

## SUPPLEMENTAL INFORMATION

Supplemental information includes Supplementary experimental procedures, Figs S1-S7 and Table ST1-ST6

## Author contributions

SAAR performed experiments, crystallization, structure determination and data analysis; LA and YAK performed experiments and contributed with reagents; AK assembled and purified ubiquitin chains; TG and DS performed the SAXS experiment and data analysis; YK supervised the research, wrote the manuscript and secured funding.

## ACKNOWLEDGEMENTS

We thank members of the Kulathu lab, specifically Drs Kwasna, Matthews and McFarland for discussions and critical comments on the manuscript, Dr. Garib Murshudov, MRC LMB for advice. Crystallographic data were collected at the European Synchrotron Radiation Facility (ESRF) beamlines ID23-1, ID23-2, ID-29, ID30B and at Diamond Light Source (DLS) beamline I03. The SAXS measurements were made at P12 beamline, PETRA III, DESY, Hamburg. TG was supported by the iNEXT project (653706) funded by the Horizon 2020 program of the European Commission. This work was supported by the Medical Research Council UK (MC_UU_00018/3), EMBO Young Investigator Programme, ERC Starting grant (677623).

## Materials Availability

Plasmids used in this study have been deposited with and will be distributed by MRC PPU reagents and services (https://mrcppureagents.dundee.ac.uk/)

## Data availability

All crystallographic data have been deposited and the PDB IDs are: MINDY1:K48-Ub2: 6TUV, MINDY2:K48-Ub2: 6Z7V, P138A C137A:K48-Ub2: 6TXB, MINDY1 Y114F: 6YJG, MINDY1 T335D: 6Y6R, MINDY1 P138A: 6Z90, MINDY2^apo^: 6Z49

The SASBDB IDs for the SAXS data in this study are: MINDY2^apo^: SASDJ93, MINDY2-Ub^Prg^: SASDJA3, MINDY2:K48-Ub2: SASDJB3, MINDY2:K48-Ub3: SASDJC3, MINDY2:K48-Ub4: SASDJD3, MINDY2:K48-Ub5: SASDJE3

