## Supplementary Information for "Mechanism of activation and regulation of Deubiquitinase activity in MINDY1 and MINDY2"

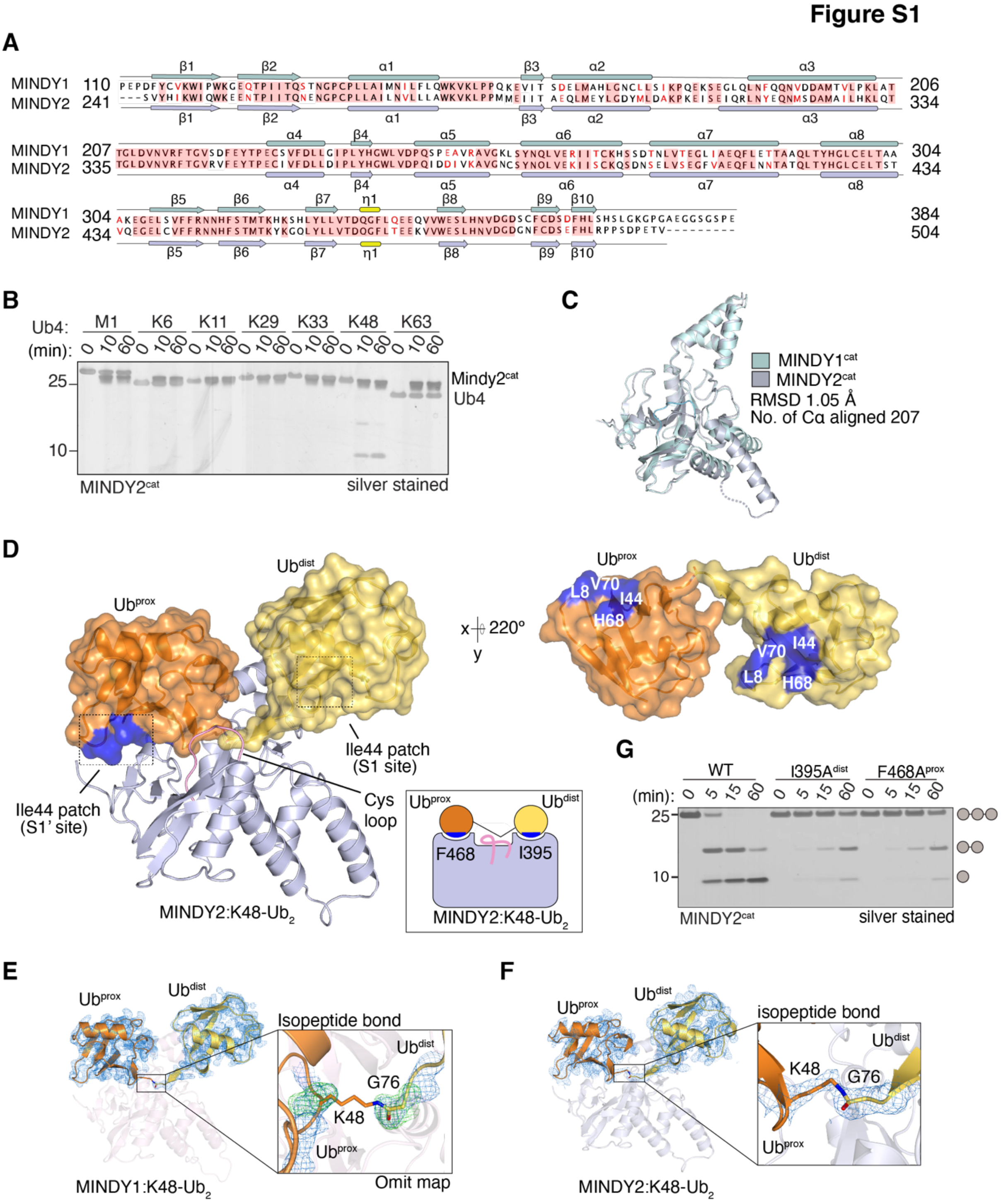

**Figure S1**

**A**) Secondary structure alignment of MINDY1 and MINDY2 based on their crystal structures using ESPRIPT webserver (http://espript.ibcp.fr/ESPript/ESPript/ ). The secondary structure elements of MINDY1 and MINDY2 are shown in pale cyan and light orange respectively.

**B)** Silver stained gels of DUB assays testing activity and specificity of polyUb cleavage by MINDY2 catalytic domain (241–504). 1.6 *μ*M of DUB was incubated with 2.2 μM of tetraUb chains for the indicated time points.

**C**) Superposition of the crystal structures of the minimal catalytic domains of MINDY1 (pale-cyan) and MINDY2 (light-orange) shown in cartoon representation.

**D**) The MINDY2:K48-diUb complex crystal structure is shown with MINDY2 in cartoon (blue white). Ub molecules are depicted with transparent surfaces (tv-orange:$Ub^{prox}$ and yelloworange: $Ub^{dist}$). I44 patches on Ub are coloured blue and an alternate view of the bound diUb rotated by 220° along the x-axis is shown on the right side. Schematic representation of MINDY2:K48-diUb complex (inset).

**E**) The crystal structure of MINDY1:K48-diUb complex with electron density for both proximal and distal ubiquitin (orange and light-brown) contoured at 1.0 σ. In the inset, the $F$o – $F$c omit electron density map was generated from coefficients calculated by removing G76 and K48 from the distal and proximal ubiquitin moieties, respectively, from the atomic model for 10 cycles of refinement. The $F$o–$F$c map is contoured at 3.0 σ whereas 2$F$o–$F$c map is contoured at 1.0 σ.

**F**) As in (E) for the MINDY2:K48-diUb complex.

**G**) DUB assay monitoring the hydrolysis of K48-linked triUb by MINDY2 and the indicated S1 and S1' site mutants.

### Figure S2

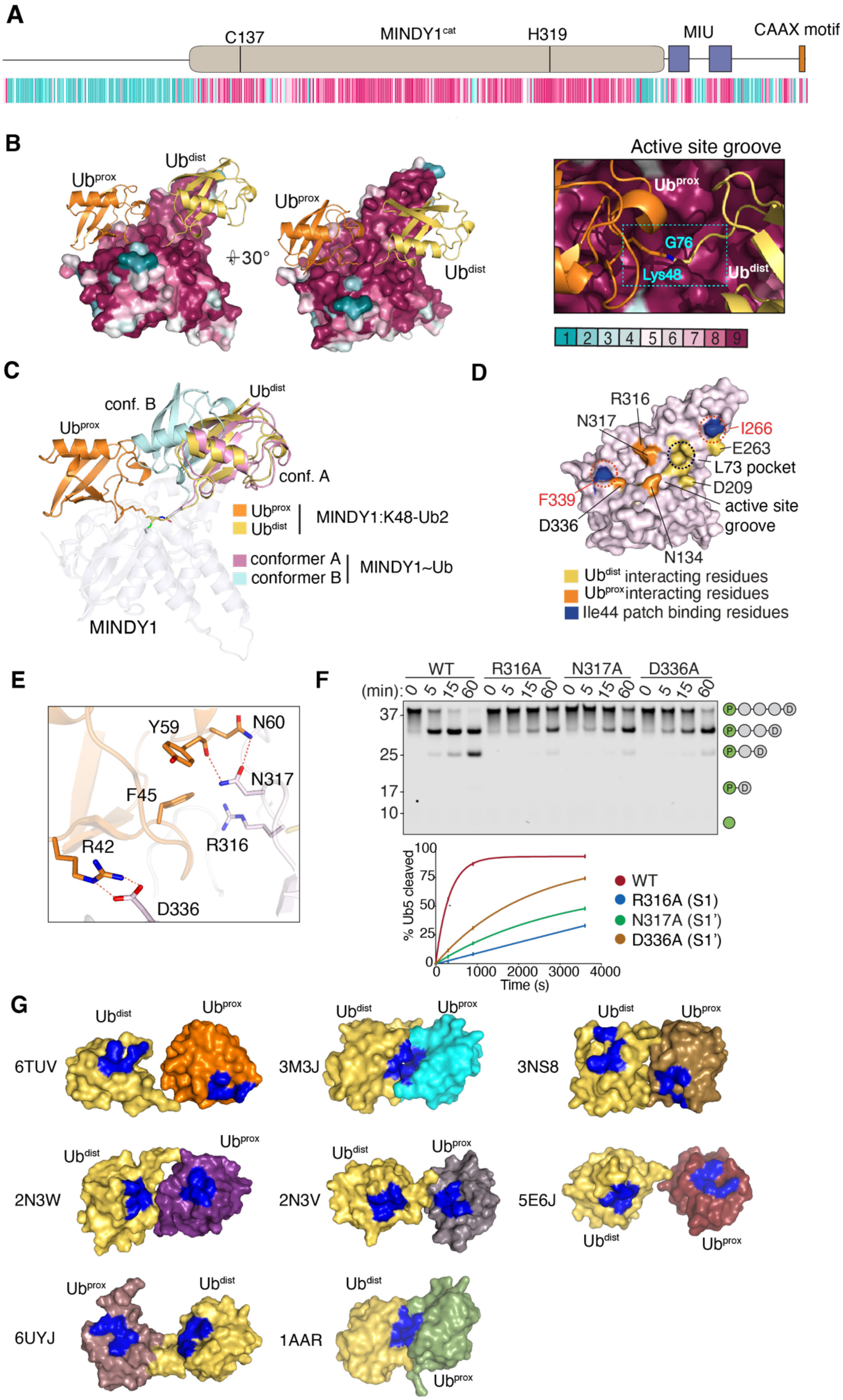
A
C137
MINDY1cat
H319
MIU
CAAX motif
B
Ubdist
Ubprox
30°
Active site groove
G76
Lys48
1 2 3 4 5 6 7 8 9
C
conf. B
conf. A
MINDY1
Ubprox Ubdist MINDY1:K48-Ub2
conformer A conformer B MINDY1~Ub
D
R316
N317
I266
E263
L73 pocket
D209
F339
active site groove
D336
N134
Ubdist interacting residues
Ubprox interacting residues
Ile44 patch binding residues
E
Y59
N60
N317
F45
R42
R316
D336
F
WT
R316A
N317A
D336A
(min): 0 5 15 60
37
25
17
10
% Ub5 cleaved
Time (s)
WT
R316A (S1)
N317A (S1')
D336A (S1')
G
6TUV
3M3J
3NS8
2N3W
2N3V
5E6J
6UYJ
1AAR

**Figure S2**

**A-B**) The evolutionary conservation score of MINDY1 was calculated using the crystal structure of MINDY1 and primary sequences from 19 species using the ConSurf webserver. The conservation score was projected both on to the primary sequence (A) and the surface representation of the crystal structure of MINDY1 (panel B) The distal and proximal ubiquitin binding region has been highlighted. In the inset, a close-up view reveals conservation of the active site and the catalytic groove (cyan dashed box) accommodating the scissile bond across the active site.

**C**) Superposition of the MINDY1:K48-Ub2 complex with MINDY1~$Ub^{Prg}$ complex reveals that proximal ubiquitin stabilises the binding of ubiquitin onto the S1 site.

**D**) Surface representation of MINDY1 highlighting key interaction interfaces.

**E)** Close-up view of the critical polar interaction between MINDY1 (pink) and the proximal ubiquitin (orange).

**F**) DUB assay comparing activity of different S1' site mutants at cleaving fluorescently-labelled pentaUb. The percent hydrolysis of K48 linked polyubiquitin chains for the different mutants is plotted against time for the DUB assay (bottom).

**G**) Comparison of different K48-linked diUb structures shown in surface representation with the I44 patches highlighted in blue. MINDY1:K48-diUb (PDB ID: 6TUV); Solution structure of the Rpn1 T1 site with K48-linked diUb in the contracted binding mode (PDB ID: 2N3W) and in the extended binding mode (PDB ID: 2N3V) (Chen *et al*, 2016); A new crystal form of K48-linked diUb (PDB ID: 3M3J) (Trempe *et al*, 2010); Crystal structure of an open conformation of K48-linked diUb at pH 7.5 (PDB ID: 3NS8) ; Crystal structure of diUb bound to SARS PLpro (PDBID: 5E6J) (Békés *et al*, 2016); hRpn13:hRpn2:K48-diUb structure (PDB ID: 6UYJ) (Lu *et al*, 2020) and crystal structure of a diUb and model for interaction with E2 (PDBID:1AAR) (Cook *et al*, 1992).

**Figure S3**

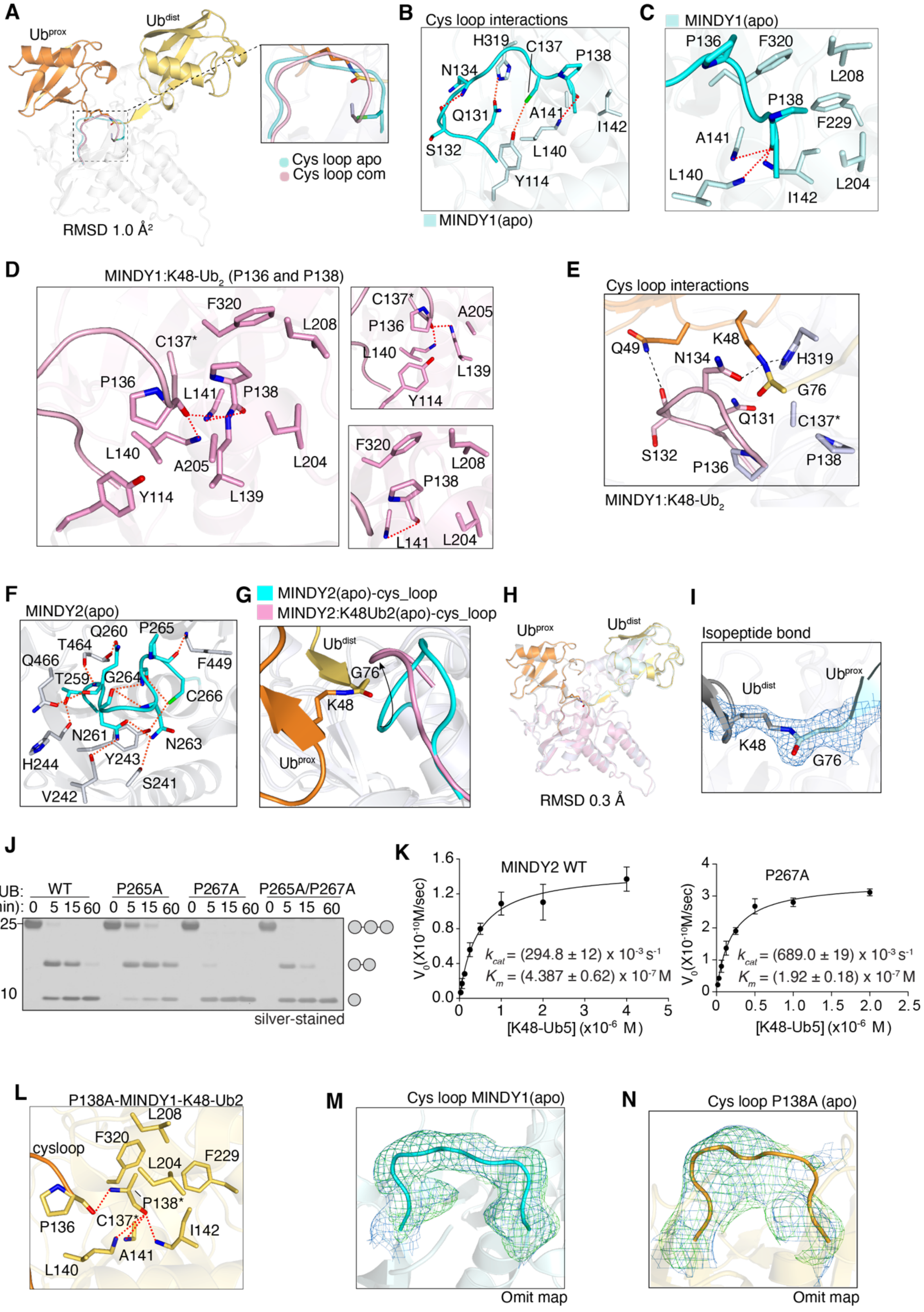

A
Ub^prox
Ub^dist
Cys loop apo
Cys loop com
RMSD 1.0 Å²
B
Cys loop interactions
H319
C137
P138
N134
Q131
A141
I142
S132
L140
Y114
MINDY1(apo)
C
MINDY1(apo)
P136
F320
L208
P138
F229
A141
L140
L204
I142
D
MINDY1:K48-Ub2 (P136 and P138)
F320
L208
C137*
P136
L141
P138
L140
A205
L204
Y114
L139
A205
L139
F320
L141
E
Cys loop interactions
Q49
K48
N134
H319
G76
Q131
C137*
S132
P136
P138
MINDY1:K48-Ub2
F
MINDY2(apo)
Q260
P265
T464
Q466
F449
T259
G264
C266
N261
N263
H244
Y243
S241
V242
G
MINDY2(apo)-cys_loop
MINDY2:K48Ub2(apo)-cys_loop
Ub^dist
G76
K48
Ub^prox
H
Ub^prox
Ub^dist
RMSD 0.3 Å
I
Isopeptide bond
Ub^dist
Ub^prox
K48
G76
J
DUB: WT P265A P267A P265A/P267A
(min): 0 5 15 60 0 5 15 60 0 5 15 60 0 5 15 60
25
10
silver-stained
K
MINDY2 WT
V0(X10^-10M/sec)
kcat = (294.8 ± 12) x 10^-3 s^-1
Km = (4.387 ± 0.62) x 10^-7 M
[K48-Ub5] (x10^-6 M)
P267A
kcat = (689.0 ± 19) x 10^-3 s^-1
Km = (1.92 ± 0.18) x 10^-7 M
[K48-Ub5] (x10^-6 M)
L
P138A-MINDY1-K48-Ub2
L208
F320
cysloop
L204
F229
P138*
P136
C137*
I142
L140
A141
M
Cys loop MINDY1(apo)
Omit map
N
Cys loop P138A (apo)
Omit map

**Figure S3**

**A**) Cartoon representation of the crystal structures of MINDY1(apo) and the MINDY1:K48-$Ub_2$ complexes (RMSD 1.0 Å). Close up view of the Cys loop in the apo state (cyan) and active state (pink).

**B**) The hydrogen bond network of the Cys loop of MINDY1(apo). Dashed lines indicate hydrogen bonds.

**C**) Hydrogen bonds formed by the backbone of P138 in MINDY1 (apo).

**D**) Close up views of hydrophobic interactions (left) and hydrogen bonding with P136 and P138 (right) in the MINDY1:K48-$Ub_2$ complex.

**E**) A close-up image of the key interactions between $Ub^{prox}$ and the Cys loop is shown.

**F**) Interaction network of the Cys loop in MINDY2 apo.

**G**) Superposition of the Cys loop in MINDY2 (apo) (dark brown) and MINDY2:K48-$Ub_2$ complex (cyan). The incoming isopeptide can be seen clashing with the Cys loop in MINDY2 apo.

**H**) Superposition of MINDY1:K48-$Ub_2$ and MINDY1 P138A:K48-$Ub_2$ complexes.

**I)** Close-up view showing electron density *2Fo-Fc* map contoured at 1.0 $\sigma$ of the isopeptide bond of the K48-$Ub_2$ from the P138A:K48-$Ub_2$ complex.

**J**) DUB assay monitoring K48-$Ub_3$ chain hydrolysis by MINDY2 WT and the mutants P265A, P267A and the double mutant P265A P267A. These two prolines flank the catalytic cysteine (C266).

**K**) Steady-state enzyme kinetics of K48-linked pentaUb cleavage by MINDY2 WT and the P267A mutant. The DUB was incubated with varying concentrations of fluorescently labelled pentaUb. (n= 2; mean ± SD).

**L**) Close-up view of the MINDY1-P138A:K48-$Ub_2$ complex showing the hydrogen bonding network of the mutant Cys loop.

**M-N**) Omit maps for $MINDY1^{apo}$ and MINDY1 P138A Cys loops. The $|F_o|-|F_c|$ electron density map was generated from coefficients calculated by deleting the Cys loop residues from the respective atomic models for 10 cycles of refinement. The $F_o-F_c$ map is contoured at 3.0 $\sigma$ and the $2F_o-F_c$ map is contoured at 1.0 $\sigma$.

**Figure S4**

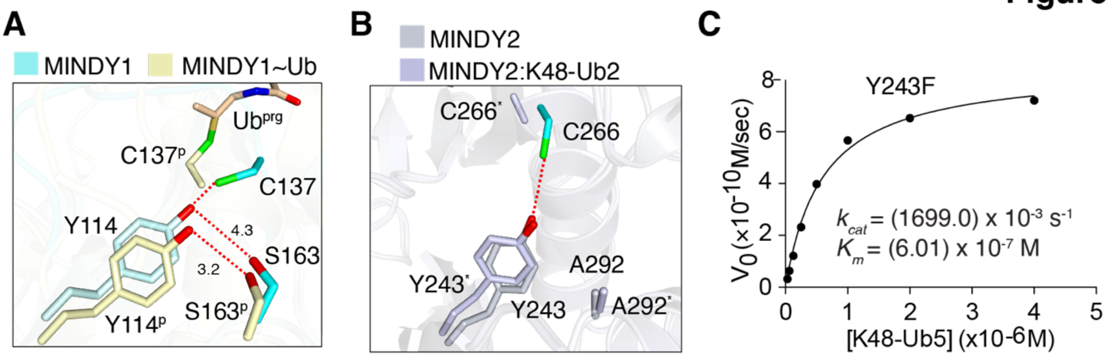

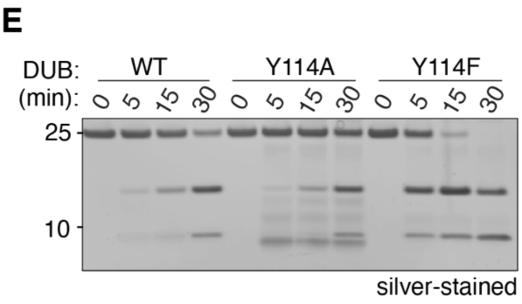

D

| Protein | $K_{cat}$ ($x10^{-3}$ $s^{-1}$) | $K_m$ ($x10^{-7}$M) |
|---|---|---|
| WT | 294.8 ± 12.0 | 4.387 ± 0.62 |
| Y243F | 1699.0 | 6.01 |

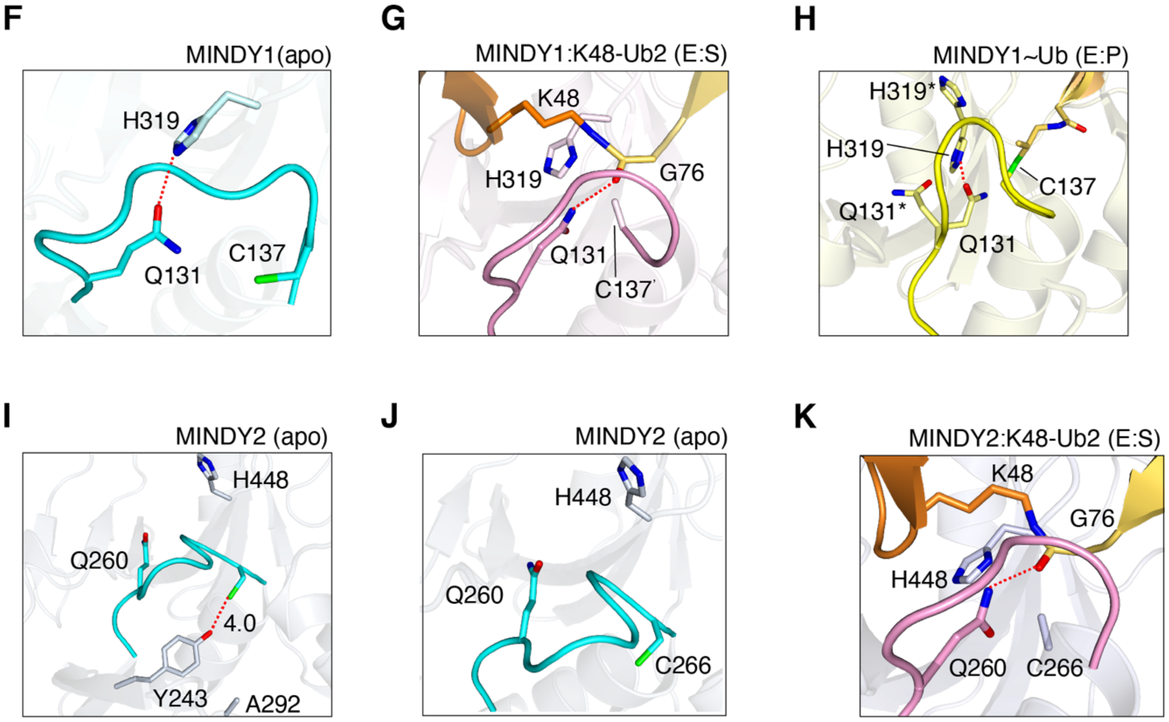

**Figure S4**

**A)** Close up view of Y114 in a superposition between MINDY1 (apo) and MINDY1~Ub$^{Prg}$. Dotted lines indicate hydrogen bond.

**B**) Close up view of superposition of MINDY2 apo and MINDY2:K48-Ub2 complex showing Y243. A292, the equivalent residue of S163 in MINDY2 does not induce lateral movement of the tyrosine observed in MINDY1. Asterisk indicates residues in complex.

**C**) Steady-state enzyme kinetics of K48-linked pentaUb cleavage by MINDY2 Y243F mutant. (n=1)

**D**) Table summarizing *kcat* and *Km* values of MINDY2 WT and MINDY2 Y243F mutant.

**E**) DUB assay monitoring the cleavage of K48linked-triUb chain by MINDY1 WT and Y114A and Y114F mutants.

**F**) A close-up view of MINDY1 apo catalytic site. The catalytic cysteine (C137) is rotated away from hydrogen bonding distance with the catalytic histidine (H319). Dotted lines indicate hydrogen bond.

**G**) A close-up view of the catalytic site in the MINDY1:K48-$Ub_2$ complex. The isopeptide bond can be seen interacting with the catalytic H319. Q131 is seen interacting with the isopeptide bond. Dotted lines indicate hydrogen bond.

**H**) A close-up view of the catalytic site in the MINDY1~UbPrg complex, where both H319 and Q131 exist in two alternate conformations. Dotted lines indicate hydrogen bond.

**I**) A close-up view of the catalytic site in MINDY2 showing key residues. The catalytic cysteine (C266) in MINDY2 is out of plane with the other catalytic residues and instead can be seen interacting with the non-catalytic Y243. Unlike MINDY1, the oxyanion forming glutamine (Q260) is not able to form any bonds with the flipped out catalytic histidine (H448). Dashed lines indicate hydrogen bond.

**J**) A close-up view of MINDY2 (apo) showing the catalytic cysteine (C266) rotated away from the hydrogen bonding distance with the catalytic histidine (H448).

**K**) View of the catalytic site in the MINDY2:K48-$Ub_2$ complex showing catalytically productive rearrangements upon K48-diUb binding. The catalytic histidine (H448) is seen to have flipped in plane with the catalytic cysteine (C266*). The oxyanion forming glutamine (Q260) is seen interacting with the isopeptide bond of the bound K48 chain.

### Figure S5

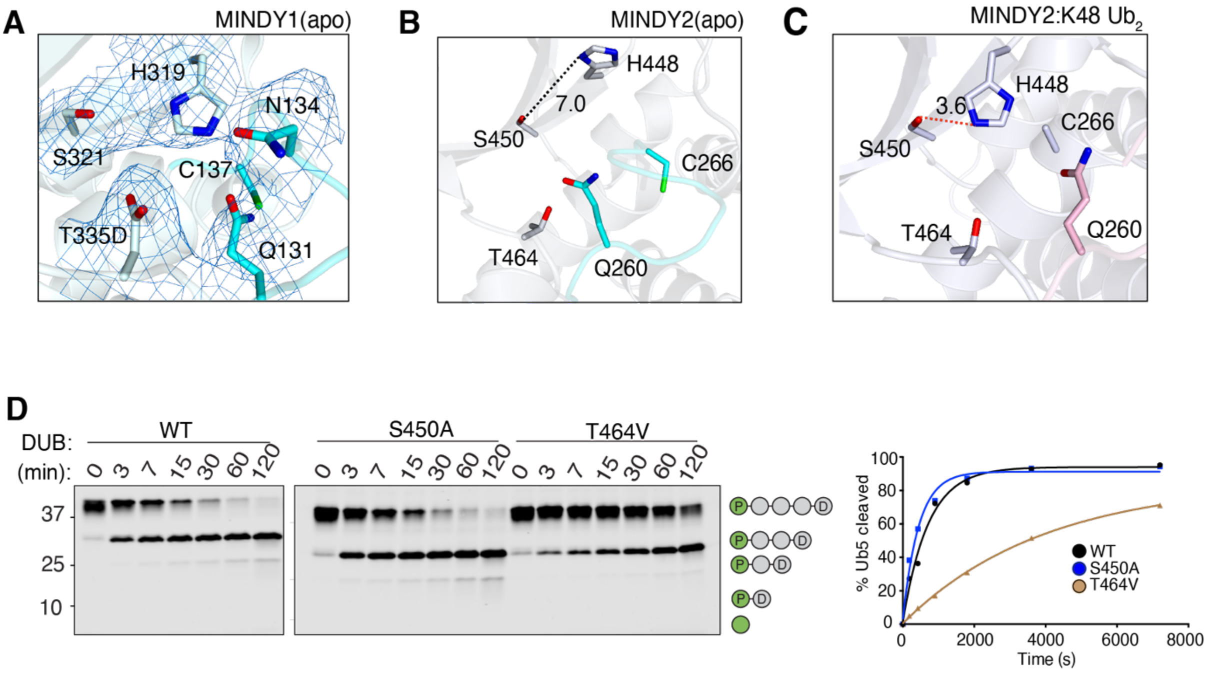

**Figure S5**

**A**) Close up view of the catalytic site in the crystal structure of MINDY1 T335D showing the formation of an ionic bond between T335D and the catalytic histidine (H319). The respective residues are shown with *2Fo-Fc* electron density contoured at 1.0 σ.

**B**)- **C**) Close up view of the catalytic site architecture in inhibited state MINDY2 apo (B) and in the active state in MINDY2:K48-$Ub_2$ (C).

**D**) DUB assay monitoring cleavage of fluorescently labelled K48-linked pentaUb chains by MINDY2 and the indicated mutants. Quantification of pentaUb cleavage is shown on the right.

### Figure S6

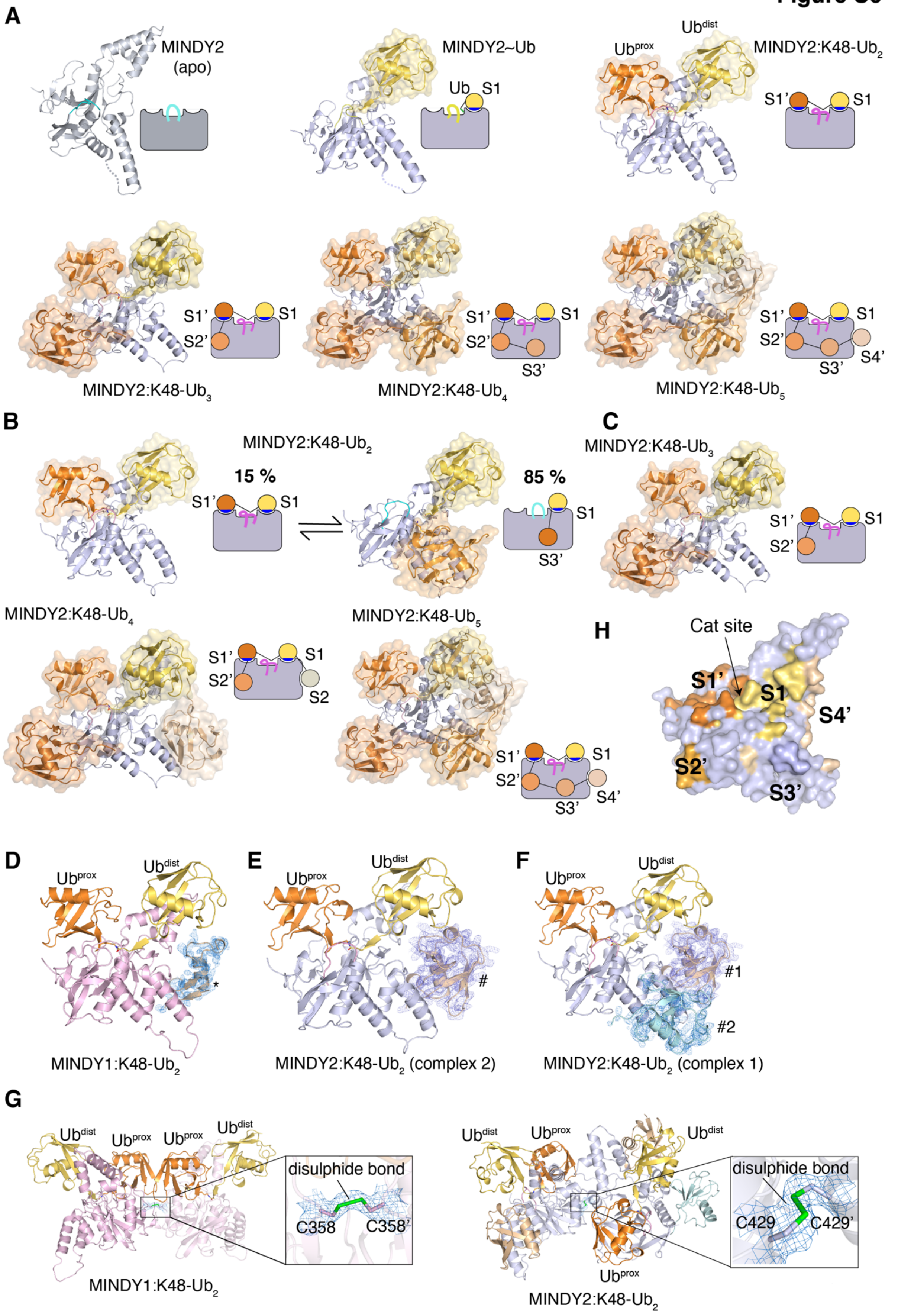

A
MINDY2 (apo)
MINDY2~Ub
Ub S1
Ub^dist
Ub^prox
MINDY2:K48-Ub2
S1' S1
S1' S1 S2'
MINDY2:K48-Ub3
S1' S1 S2' S3'
MINDY2:K48-Ub4
S1' S1 S2' S3' S4'
MINDY2:K48-Ub5
B
MINDY2:K48-Ub2
15 %
85 %
S1' S1
S1 S3'
C
MINDY2:K48-Ub3
S1' S1 S2'
MINDY2:K48-Ub4
S1' S1 S2' S2
MINDY2:K48-Ub5
S1' S1 S2' S3' S4'
H
Cat site
S1' S1 S4' S2' S3'
D
Ub^prox Ub^dist
*
MINDY1:K48-Ub2
E
Ub^prox Ub^dist
#
MINDY2:K48-Ub2 (complex 2)
F
Ub^prox Ub^dist
#1
#2
MINDY2:K48-Ub2 (complex 1)
G
Ub^dist Ub^prox Ub^prox Ub^dist
disulphide bond
C358 C358'
MINDY1:K48-Ub2
Ub^dist Ub^prox Ub^dist
Ub^prox
disulphide bond
C429 C429'
MINDY2:K48-Ub2

**Figure S6**

**A**) The models used in the curve fitting with the SAXS data are depicted along with a schematic representation. The MINDY2:K48-Ub4 model was a very poor fit and was not used further.

**B)** Models validated with considerable agreement.

**C)** Models observed and validated in SAXS measurements with some deviations and improved curve fitting post normal mode analysis run.

**D**) Additional electron density for a partial ubiquitin molecule present near the S1 site in the MINDY1:K48-$Ub_2$ complex.

**E**) Additional electron density for a full ubiquitin molecule present near the S1 site of MINDY2:K48-diUb complex.

**F**) Additional electron density for a fourth ubiquitin bound at a helix-loop-helix region of the MINDY2:K48-$Ub_2$ complex.

**G**) Disulphide bond formed between DUB in the ASU with DUB in symmetry-related molecule is highlighted in the inset for MINDY1 C358-C358 (left) and MINDY2 C429-C429 (right). Electron density *2Fo-Fc* contoured at 1$\sigma$ for the cysteines forming the disulphide bond (inset).

**H**) Location of Ub binding sites is shown on the surface of MINDY2

### Figure S7

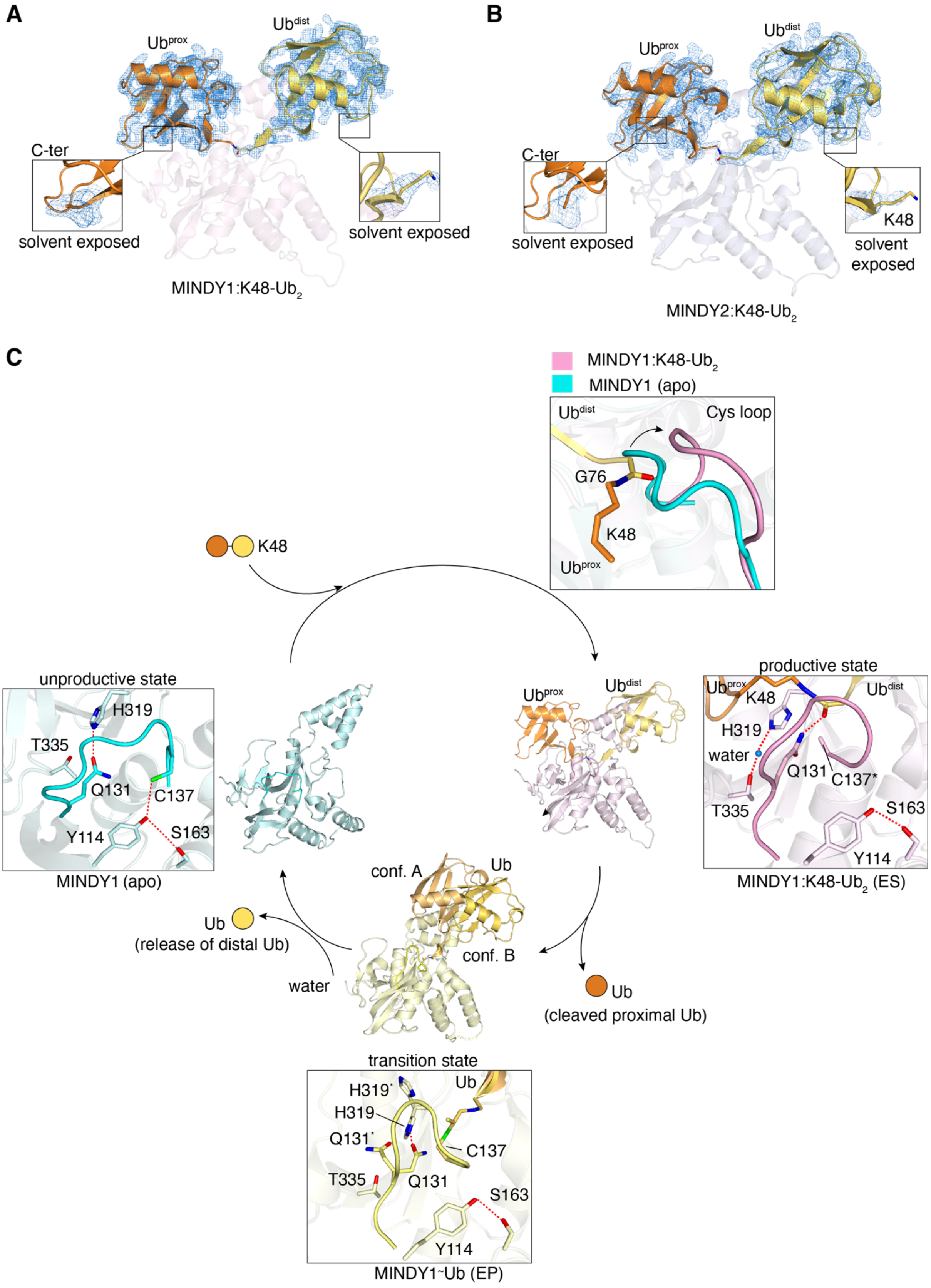
A
Ub$^{prox}$
Ub$^{dist}$
C-ter
solvent exposed
solvent exposed
MINDY1:K48-Ub$_2$
B
Ub$^{prox}$
Ub$^{dist}$
C-ter
solvent exposed
K48
solvent exposed
MINDY2:K48-Ub$_2$
C
MINDY1:K48-Ub$_2$
MINDY1 (apo)
Ub$^{dist}$
Cys loop
G76
K48
Ub$^{prox}$
K48
unproductive state
H319
T335
Q131
C137
Y114
S163
MINDY1 (apo)
Ub$^{prox}$
Ub$^{dist}$
productive state
Ub$^{prox}$
K48
Ub$^{dist}$
H319
water
Q131
C137*
T335
S163
Y114
MINDY1:K48-Ub$_2$ (ES)
conf. A
Ub
conf. B
Ub
(release of distal Ub)
water
Ub
(cleaved proximal Ub)
transition state
H319'
Ub
H319
Q131'
C137
T335
Q131
S163
Y114
MINDY1~Ub (EP)

**Figure S7**

**A)** Crystal structure of MINDY1:K48-diUb complex with *2Fo-Fc* electron density for both proximal and distal ubiquitin (tv-orange and yellow-orange) contoured at 1.0 σ. In the inset (right side), the solvent exposed C-terminus of proximal ubiquitin (contoured at 1.0 σ) is shown whereas on the right side solvent exposed K48 of distal ubiquitin is shown (contoured at 0.5 σ).

**B)** Structure of MINDY2:K48-diUb complex with *2Fo-Fc* electron density for both proximal and distal ubiquitin (light-orange and yellow-orange) contoured at 1.0 σ. In the inset, the solvent exposed C-terminus of $Ub^{prox}$ (contoured at 1.0 σ) and K48 of $Ub^{dist}$ (contoured at 0.8 σ) are shown.

**C)** Related to Fig 7, this shows the structures and the transitions in the catalytic cycle of MINDY1 going from inactive state to substrate bound active state and MINDY1 in the transition state bound to product intermediate.

**Table S1: Summary of major ubiquitin interactions with DUBs**

| PDB ID | Title of Structure | Ub$^{prox}$ | Ub$^{dist}$ |
|---|---|---|---|
| 6TUV, 6Z7V | MINDY1 in complex with K48-linked diubiquitin | **L 8, I44**<br>**V70, F45**<br>R42 | **L 8, I44**<br>**V70, L73**<br>R42 |
| 6NJD | Crystal structure of RavD from Legionella pneumophila complexed with Met-1 linked di-ubiquitin | H61, K33<br>G35, Q2<br>E16, E64<br>D32 | L8, I44, V70, L73,<br>L73, R74<br>K6, R42, K48, R72 |
| 5LRV | Structure of Cezanne/OTUD7B OTU domain bound to K11-linked diubiquitin | K33, D32<br>E34, G35<br>K33, D32 | I44, I36, L71, L73,<br>Ile13, I30<br>L69, L8, R42, R72,<br>R74 |
| 2ZNV | Crystal structure of human AMSH-LP DUB domain in complex with K63-linked ubiquitin dimer | F4, Gln62<br>E64, Asn60<br>K63 | L73, V70, I44, L8,<br>I36, L71<br>R42, R72, R74, K48,<br>K6, His68<br>D39, R74 |
| 4NQL | The crystal structure of the DUB domain of AMSH orthologue, Sst2 from S. pombe, in complex with lysine 63-linked diubiquitin | F4, Gln62<br>E64, Q2<br>K63 | L73, I44, V70, L8,<br>L71, I36<br>L69, R42<br>R72, His68 |
| 5OHP | Crystal structure of USP30 (C77A) in complex with K6-linked diubiquitin | L8, F4<br>K6, His68<br>E64, K11 | L71, L73, I36, Pro37,<br>F4, G76<br>G75, R74, R42, R72,<br>K48, K6<br>E16, F4, K6 |
| 3WXE | Crystal structure of CYLD USP domain (C596S) in complex with M1-linked diubiquitin | F4, M1<br>Q2, Thr12<br>E64, Glu18<br>K6, E16<br>F4, K63 | V70, L8<br>I36, L69<br>L71, L73<br>G75, R74<br>R42, R72<br>Glu51, K6 |
| 3WXG | Crystal structure of CYLD USP domain (C596A) in complex with K63-linked diubiquitin | F4, E64, K63, Glu18<br>K29, K6, E16 | V70, L8, I36, L69,<br>L71, L73<br>G75, G76, R74, R42,<br>R72, Glu51<br>K6 |
| 3ZNZ<br><br><br><br>5KSL<br><br><br>5OE7 | Crystal structure of OTULIN OTU domain (C129A) in complex with M1- di ubiquitin<br>Structure of OTULIN bound to the M1-linked diubiquitin activity probe<br>Gumby/Fam105B in complex with linear di-ubiquitin | K33, D32<br>M1, Q2<br>E34, K63<br>K33, E34<br>D32, E64<br>K63, K29<br>E16, K33 | L8, I44, V70<br>G75, G76<br>R42, Glu32<br>K11 |

**Colour coding:** Hydrophobic; Hydrogen bond; Ionic bond; Cation-Pi

#### Table S2. Data collection and refinement statistics

| | MINDY1$^{Y11F}$ | MINDY1$^{T335D}$ |
|---|---|---|
| **Data Collection** | | |
| Beamline | ID23-2, ESRF | I03, DLS |
| Wavelength (Å) | 0.87313 | 0.97628 |
| Space group | $P4_122$ | $P4_122$ |
| Total reflections | 378950 (74980) | 70094 (11564) |
| a, b, c (Å) | 102.16, 102.16, 162.71 | 99.81, 99.81, 164.11 |
| α, β, γ (°) | 90.00, 90.00, 90.00 | 90.00, 90.00, 90.00 |
| Resolution (Å) | 72.240-3.28 | 85.28-3.32 |
| $R_{merge}$ | 0.278 (1.734) | 0.057 (1.022) |
| I/σ(I) | 11.5 (2.5) | 13.2 (1.2) |
| Completeness (%) | 100.0 (99.8) | 99.4 (97.4) |
| Multiplicity | 27.3 (27.0) | 5.5 (4.6) |
| CC1/2 | 0.998 (0.902) | 0.999 (0.721) |
| **Refinement** | | |
| Resolution (Å) | 72.2109-3.28 | 85.21-3.32 |
| $R_{work/}$ $R_{free}$ | 0.205/0.246 | 0.20/0.24 |
| **No. of Atoms** | | |
| Protein | 1984 | 1993 |
| Ligand | N/A | N/A |
| Water | N/A | N/A |
| **B-Factors (Å$^2$)** | | |
| Protein | 93.47 | 168.15 |
| Ligand | N/A | N/A |
| Water | N/A | N/A |
| **RMSDs** | | |
| Bond length (Å) | 0.010 | 0.001 |
| Bond angles (°) | 2.040 | 1.792 |
| **Ramachandran Plot (%)** | | |
| Favored region | 90 | 89 |
| Allowed region | 10 | 11 |
| Outlier region | 0 | 0 |
| PDB ID | **6YJG** | **6Y6R** |

DLS, Diamond Light Source; ESRF, European Synchrotron Radiation Facility; RMSD, root-mean-square deviation. Values for the highest-resolution shell are shown in parentheses

#### Table S3. Data collection and refinement statistics

| | MINDY2 | MINDY2:K48-$Ub_2$ |
|---|---|---|
| **Data Collection** | | |
| Beamline | ID29, ESRF | ID29, ESRF |
| Wavelength (Å) | 1.07252 | 0.97625 |
| Space group | $P12_11$ | $P2_12_12_1$ |
| Total reflections | 344672 (18610) | 176139 (20372) |
| a, b, c (Å) | 67.11, 120.42, 78.38 | 94.01, 117.60, 125.88 |
| α, β, γ (°) | 90.00, 92.94, 90.00 | 90.00, 90.00, 90.00 |
| Resolution (Å) | 47.72-2.00 | 47.79-2.65 |
| $R_{merge}$ | 0.045 (0.599) | 0.072 (0.544) |
| I/σ(I) | 13.9 (2.1) | 11.8 (2.1) |
| Completeness (%) | 99.7 (99.7) | 98.40 (99.60) |
| Multiplicity | 4.1 (4.1) | 4.4 (4.5) |
| CC1/2 | 0.999 (0.814) | 0.998 (0.704) |
| **Refinement** | | |
| Resolution (Å) | 47.72-2.00 | 47.75-2.65 |
| $R_{work}$/ $R_{free}$ | 0.215/0.251 | 0.22/0.26 |
| **No. of Atoms** | | |
| Protein | 7794 | 7686 |
| Ligand | 42 | 43 |
| Water | 115 | 39 |
| **B-Factors (Å$^2$)** | | |
| Protein | 47.93 | 69.20 |
| Ligand | 65.89 | 86.30 |
| Water | 38.86 | 52.99 |
| **RMSDs** | | |
| Bond length (Å) | 0.008 | 0.007 |
| Bond angles (°) | 1.567 | 1.577 |
| **Ramachandran Plot (%)** | | |
| Favored region | 97.0 | 98.0 |
| Allowed region | 3.0 | 2.0 |
| Outlier region | 0 | 0 |
| PDB ID | **6Z49** | **6Z7V** |

DLS, Diamond Light Source; ESRF, European Synchrotron Radiation Facility; RMSD, root-mean-square deviation. Values for the highest-resolution shell are shown in parentheses

#### Table S4: SAXS data collection

| | | | | | | |
|---|---|---|---|---|---|---|
| **Data collection parameters** | | | | | | |
| Radiation source | PETRA III (DESY, Hamburg, Germany) | | | | | |
| Beamline | EMBL P12 | | | | | |
| Detector | PILATUS 6M | | | | | |
| Beam geometry (mm, FWHM) | 0.12 x 0.20 | | | | | |
| Wavelength (nm) | 0.12 | | | | | |
| Sample-detector distance (m) | 3.0 | | | | | |
| Momentum transfer *s* range ($nm^{-1}$) | 0.01 – 7.0 | | | | | |
| Exposure time (s) | 1 s (SEC-SAXS mode) | | | | | |
| Temperature (°C) | 20 | | | | | |
| Buffer | 20 mM Hepes, 100 mM NaCl, 5 mM DTT, pH 7.5 | | | | | |
| column | Superdex 200 10/300 | | | | | |
| **Overall Parameters** | | | | | | |
| | MINDY 2 Apo | MINDY 2 Ub 1 | MINDY 2 Ub 2 | MINDY 2 Ub 3 | MINDY 2 Ub 4 | MINDY 2 Ub 5 |
| Number of ubiquitins | 0 | 1 | 2 | 3 | 4 | 5 |
| Concentration (mg/ml) | 8.8 | 10.5 | 9.9 | 12.2 | 13.7 | 17.1 |
| $R_g$ from Guinier approximation (nm) | 2.1±0.1 | 2.3±0.1 | 2.5±0.1 | 2.8±0.2 | 2.9±0.2 | 2.8±0.2 |
| $D_{max}$ (nm) | 6.5±0.3 | 6.8±0.3 | 8.0±0.4 | 8.8±0.5 | 9.2±0.5 | 9.0±0.6 |
| Excluded (Porod) volume ($nm^3$) | 45±4 | 58±6 | 68±7 | 97±9 | 111±10 | 116±10 |
| Molecular weight from Porod invariant, (kDa) | 28±3 | 35±4 | 43±5 | 60±6 | 69±7 | 73±7 |
| Molecular weight from sequence (kDa) | 31.0 | 39.5 | 48.0 | 56.5 | 65.0 | 73.5 |
| **Data analysis and modelling** | | | | | | |
| Primary data reduction | SASFLOW | | | | | |
| Data processing | PRIMUS/CHROMIXS | | | | | |
| Calculation of theoretical data | CRYSOL | | | | | |
| $\chi^2$ Crysol | 2.04 | 1.47 | 5.23 | 2.07 | 1.65 | 3.33 |
| Model refinement | SREFLEX | | | | | |
| $\chi^2$ SREFLEX | 1.49 | - | - | 1.15 | - | 1.54 |
| RMSD | 3.91 | - | - | 5.01 | - | 2.12 |
| Multicomponent Mixture Analysis | OLIGOMER | | | | | |
| $\chi^2$ OLIGOMER | - | - | 1.90 | - | - | - |
| SASBDB accession code | SASDJ93 | SASDJA3 | SASDJB3 | SASDJC3 | SASDJD3 | SASDJE3 |

#### Table S5: key resources table

| Reagent or Resource | Source | Additional information |
|---|---|---|
| Bacterial Strains | | |
| BL21(DE3) cells | New England Biolabs | Cat# C2527H |
| Chemicals, Peptides, and Recombinant Proteins | | |
| Propargylamine | Sigma Aldrich | Cat# P50900-5G |
| Gluthahione Sepharose 4B | Expedeon | Cat# AGSCUST |
| **Deposited Data** | | |
| Structure factor and coordinates files | RCSB-PDB | https://www.rcsb.org/ |
| **Software and Algorithms** | | |
| Prism | Graphpad | https://www.graphpad.com/scientific-software/prism/ |
| XDS | Kabsch, 2010 | http://xds.mpimf-heidelberg.mpg.de/ |
| AIMLESS | Evans and Murshudov, 2013 | http://www.ccp4.ac.uk/html/aimless.html |
| CCP4 interface version 7.0.04 | Winn et al., 2011 | http://www.ccp4.ac.uk/ |
| Phenix | Liebschner et al., 2019 | https://www.phenix-online.org/ |
| COOT | Emsley et al., 2010 | http://www2.mrc-lmb.cam.ac.uk/personal/pemsley/coot/ |
| REFMAC5 | Murshudov et al., 1997 | http://www.ccp4.ac.uk/html/refmac5/description.html |
| PDB-REDO | Joosten et al., 2014 | http://www.cmbi.ru.nl/pdb_redo/ |
| PyMOL | | https://pymol.org/2/ |
| Adobe Illustrator | | https://www.adobe.com/uk/products/illustrator.html |

**Table S6: Details of cDNA constructs used in study**

| Protein | Expressed protein | Tag Cleaved | Vector type | Plasmid | DU number |
|---|---|---|---|---|---|
| MINDY1$^{FL}$ | GST-MINDY1 1-469 | Yes | Bacterial | pGEX6P1 | 49563 |
| MINDY2$^{FL}$ | GST-MINDY2 1-621 | Yes | Bacterial | pGEX6P1 | 46765 |
| MINDY1$^{cat}$ | GST-MINDY1 110-384 | Yes | Bacterial | pGEX6P1 | 47257 |
| MINDY2$^{cat}$ | GST-MINDY2 241-384 | Yes | Bacterial | pGEX6P1 | 53390 |
| MINDY1$^{cat}$ C137A | GST-MINDY1-C137A 110-384 | Yes | Bacterial | pGEX6P1 | 47419 |
| MINDY2$^{cat}$ C266A | GST-MINDY2 C266A 241-504 | Yes | Bacterial | pGEX6P1 | 55471 |
| I266A | GST-MINDY1-I266A 110-384 | Yes | Bacterial | pGEX6P1 | 47820 |
| F339A | GST-MINDY1-F339A 110-384 | Yes | Bacterial | pGEX6P1 | 47665 |
| I266A-F339A | GST-MINDY1-I266A F339A 110-384 | Yes | Bacterial | pGEX6P1 | 67842 |
| I395A | GST-MINDY2 I395A 241-384 | Yes | Bacterial | pGEX6P1 | 55941 |
| F468A | GST-MINDY2 F468A 241-504 | Yes | Bacterial | pGEX6P1 | 55878 |
| R316A | GST-MINDY1-R316A 110-384 | Yes | Bacterial | pGEX6P1 | 47668 |
| N317A | GST-MINDY1-N317A 110-384 | Yes | Bacterial | pGEX6P1 | 59172 |
| D336A | GST-MINDY1-D336A 110-384 | Yes | Bacterial | pGEX6P1 | 47667 |
| P138A | GST-MINDY1-P138A 110-384 | Yes | Bacterial | pGEX6P1 | 47661 |
| P138G | GST-MINDY1-P138G 110-384 | Yes | Bacterial | pGEX6P1 | 59586 |
| P138L | GST-MINDY1-P138L 110-384 | Yes | Bacterial | pGEX6P1 | 59683 |
| P138W | GST-MINDY1-P138W 110-384 | Yes | Bacterial | pGEX6P1 | 59684 |
| P136A | GST-MINDY1-P136A 110-384 | Yes | Bacterial | pGEX6P1 | 47821 |
| P136A P138A | GST-MINDY1-P136A P138A 110-384 | Yes | Bacterial | pGEX6P1 | 47815 |
| Y114A | GST-MINDY1-Y114A 110-384 | Yes | Bacterial | pGEX6P1 | 47659 |
| Y114F | GST-MINDY1-Y114F 110-384 | Yes | Bacterial | pGEX6P1 | 47671 |
| N134A | GST-MINDY1-N134A 110-384 | Yes | Bacterial | pGEX6P1 | 59152 |
| S321A | GST-MINDY1-S321A 110-384 | Yes | Bacterial | pGEX6P1 | 47670 |
| S321D | GST-MINDY1-S321D 110-384 | Yes | Bacterial | pGEX6P1 | 59666 |
| T335V | GST-MINDY1-T335V 110-384 | Yes | Bacterial | pGEX6P1 | 47757 |
| T335D | GST-MINDY1-T335D 110-384 | Yes | Bacterial | pGEX6P1 | 58884 |
| P265A | GST-MINDY2 P265A 241-504 | Yes | Bacterial | pGEX6P1 | 59611 |
| P267A | GST-MINDY2 P267A 241-504 | Yes | Bacterial | pGEX6P1 | 55477 |

| | | | | | |
|---|---|---|---|---|---|
| P265A P267A | GST-MINDY2 P265A P267A 241-384 | Yes | Bacterial | pGEX6P1 | 59610 |
| P138A C137A | GST-MINDY1C137A P138A 110-384 | Yes | Bacterial | pGEX6P1 | 55726 |
| S450A | GST-MINDY2-S450A 241-384 | Yes | Bacterial | pGEX6P1 | 55826 |
| T464V | GST-MINDY2-T464V 241-384 | Yes | Bacterial | pGEX6P1 | 55825 |
| Ubiquitin 1-76 | Ubiquitin (expressed tagless) | Tagless | Bacterial | pET24 | 20027 |
| Ubiquitin 1-75 | Ub-Intein-CBD 1-75 | Yes | Bacterial | pTXB1 | 24149 |

#### METHOD DETAILS

##### Plasmids

All cDNA constructs used in this study were generated by the Cloning team of the MRC reagents and services facility, MRC Protein Phosphorylation and Ubiquitylation Unit, University of Dundee, United Kingdom (see **table S6**).

##### Protein expression and purification

All recombinant GST-fusion proteins were expressed in E. coli strain BL21(DE3). The bacterial cell cultures were grown in 2xTY media containing 100 $\mu$g/ml ampicillin to an $OD_{600}$ of 0.6-0.8 at 37ºC. Protein expression was induced with 300 $\mu$M IPTG followed by overnight shaking at 18 °C. Cells were harvested at 4000 rpm for 15 minutes and the pellets were resuspended in GST-Lysis Buffer (50 mM Tris-HCl pH 7.5, 300 mM NaCl, 10% glycerol, 0.075% 2-mercaptoethanol, 1 mM benzamidine, 1 mM AEBSF, and complete protease inhibitor cocktail (Roche)). The resuspended cells were lysed by sonication and clarified by centrifugation at 30,000 x g for 45 min at 4ºC and the lysates were incubated with Glutathione Sepharose 4B resin (Expedion) for 2 hrs at 4 °C on a rolling shaker. Resin was washed extensively, first with high salt buffer (25 mM Tris pH 7.5, 500 mM NaCl, and 10 mM DTT) and then with low salt buffer (50 mM Tris-HCl pH 7.5, 150 mM NaCl, 10% glycerol, and 1 mM DTT). The GST tag was removed by on column cleavage with 3C protease in an overnight incubation at 4 °C. All purified proteins used for DUB assays or enzymes kinetics were quantified using nanodrop at $A_{280}$ and aliquots were flashed frozen in liquid nitrogen and stored at -80 ºC. Proteins meant for ITC, SAXS or crystallization were further purified by anion exchange chromatography (Resource Q, GE Healthcare Life Sciences) and eluted in a gradient with buffer Q (50 mM Tris-HCl pH 8.5, 1 M NaCl and 2 mM DTT), followed by size exclusion chromatography (Superdex 75 16/60, GE Healthcare Life Sciences) in a relevant buffer. Buffer I used for ITC (50mM Tris-HCl pH 7.5, 150mM NaCl and 250$\mu$M TCEP), buffer S for SAXS (20 mM HEPES pH 7.5, 100 mM NaCl, 5mM DTT) and buffer X for crystallization (50 mM Tris-HCl, 150 mM NaCl, 10 mM DTT). The purified proteins were concentrated, quantified using nanodrop and flash frozen in liquid nitrogen and stored at -80 °C.

##### Deubiquitylation assays (qualitative)

DUBs were diluted in 50 mM Tris-HCl pH 7.5, 50 mM NaCl, 10 mM DTT and incubated at room temperature (24 °C) for 10 min to fully reduce the catalytic Cys. DUB assays were subsequently carried out where 1.9 $\mu$M of K48-Ub2 or K48-Ub3 were incubated with 1.6 $\mu$M of MINDY1 in 50 mM Tris-HCl pH 7.5, 50 mM NaCl, 10 mM DTT in a reaction volume of 10 $\mu$l. For DUB assay against different linkage types, 1.9 $\mu$M of diUb or 2.2 $\mu$M of tetraUb of specific linkage types were incubated with 1.6 $\mu$M of MINDY1 or 1.6 $\mu$M of MINDY2 in 50 mM Tris-HCl pH 7.5, 50 mM NaCl, 10 mM DTT in a reaction volume of 10 μl (Figs S1B and 2H). For DUB assays comparing activity of MINDY1$^{FL}$ and MINDY1$^{cat}$ at cleaving long K48 chains, 3.5 $\mu$g of K48-Ub5-n or 2.2 $\mu$M of K48-Ub6 were incubated with 1.6 $\mu$M of MINDY1 in 50 mM Tris-HCl pH 7.5, 50 mM NaCl, 10 mM DTT in a reaction volume of 10 $\mu$l. For DUB assays comparing activity of MINDY2$^{FL}$ and MINDY2$^{cat}$ at cleaving K48 chains, 3.5 $\mu$g of K48-Ub5-n or 2.2 $\mu$M of K48-Ub6 were incubated with 0.1 $\mu$M of MINDY2 in 50 mM Tris-HCl pH 7.5, 50 mM NaCl, 10 mM DTT in a reaction volume of 10 $\mu$l.  All reactions were incubated at 30 °C and stopped at indicated time points by adding LDS buffer. The samples were separated on 4-12% SDS-PAGE gel (Life Technology) and silver stained using Pierce Silver stain kit (Thermo Fisher).

**Deubiquitylation assays (quantitative)**

K48-linked Ub chains were fluorescently labelled on the distal Ub as described previously (Abdul Rehman *et al*, 2016). Both DUBs and fluorescently labelled K48-polyUb chains were diluted in 50 mM Tris-HCl pH 7.5, 50 mM NaCl, 10 mM DTT, and 0.25 mg/ml BSA. DUBs were activated by incubation at room temperature for 10 min. The reaction mixtures containing 1 $\mu$M DUB and 500 nM fluorescently labelled K48polyUb chains were incubated at 30 °C and stopped at indicated time points by adding LDS buffer. At the indicated time points, 2.5 $\mu$l of the samples was transferred to 7.5 $\mu$l LDS sample buffer to quench the reaction. The SDS gels of these DUB assays were scanned with Odyssey® CLx Imaging System at 800 nm channel and quantified with Image Studio™ Lite software. Data from two independent experiments were fitted using nonlinear regression. Data fitting was performed using GraphPad Prism 8 software.

**Enzyme kinetics**

Steady-state kinetics of K48 linked Ub5 fluorescent chain (IR-K48-Ub5) hydrolysis by MINDY1, MINDY2 and their mutants (Y114A$^{MINDY1}$, P138A$^{MINDY1}$, Y243A$^{MINDY2}$ and P267A$^{MINDY2}$). The catalytic domain of MINDY1 and MINDY2 and their mutants were incubated with varying concentrations of K48 linked Ub5 fluorescent chains and the formation of K48 Ub4 at the early time points was quantified to obtain the initial velocities. Both DUBs and fluorescently labelled K48-polyUb chains were diluted in 50 mM Tris-HCl pH 7.5, 50 mM NaCl, 10 mM DTT, and 0.25 mg/ml BSA. At the indicated time points (0, 3, 6, 9 and 12 minutes), 2.5 $\mu$l of the samples was transferred to 7.5 $\mu$l LDS sample buffer to quench the reaction. The SDS gels of DUB assays were scanned with Odyssey® CLx Imaging System at 800 nm channel and quantified with Image Studio™ Lite software. The amount of K48-Ub4 formed was plotted against time and the data was fitted to a linear regression curve, where slope is the initial velocity, $V_o$ (M.s-1). The initial velocities obtained were plotted against the of K48 linked Ub5 fluorescent chains (substrate) concentration and the curves were fitted to Michaelis-Menten equation to estimate the $K_{cat}$ and $K_m$ (n = 2; mean ± SD). Data fitting was carried out using GraphPad Prism 8 software.

**ITC measurements**

ITC measurements were performed on MicroCal PEAQ-ITC (Malvern) at 25ºC. Prior to measurements, all proteins were dialysed into a buffer containing 50mM Tris-HCl pH 7.5, 150mM NaCl and 250$\mu$M TCEP. The syringe contained MINDY1 and was titrated into the cell which contained K48-linked polyubiquitin chains (K48-diUb, triUb or tetraUb or pentaUb). 2 $\mu$l of MINDY1 was dispensed in 4-sec duration with 130-sec spacing in between injections for a total of 16 injections. Data were analysed and titration curves were fitted using MicroCal PEAQ-ITC (Malvern) analysis software (n = 2; mean ± SD).

**Crystallization and Structure determination**

**MINDY1:K48-Ub2**

The MINDY1-catalytic domain C137A mutant construct (residues 110-384) in 50 mM Tris-HCl pH 7.5, 150 mM NaCl, and 10 mM DTT was mixed with K48-linked diUb in a 1:1 ratio and concentrated to a final concentration of 11.5 mg/ml. The crystals were grown in hanging drop 24 well plates against the well solution of 0.1 M Tris-HCl pH

8.5, 0.2 M Lithium sulphate monohydrate and 30% PEG 400. The crystals were flash frozen in cryo-protectant containing 0.1 M Tris-HCl pH 8.5, 0.2 M Lithium sulphate monohydrate and 35% PEG 400. Diffraction data were collected at ID23-1 beamline, ESRF, France (wavelength 0.9397 Å). The data sets were processed using XDS (Kabsch, 2010) and then scaled using AIMLESS (Evans and Murshudov, 2013). The structure of the complex MINDY1:K48-Ub2 was solved by molecular replacement (Phaser) using MINDY1$^{apo}$ (PDB ID: 5JKN) and Ub (PDB ID: 1UBQ) as search models. The partially built model obtained was further manually built in COOT. The complete model was obtained after iterative building and refinement with COOT (Emsley et al., 2010) and REFAMC5 (Murshudov et al., 1997). The final structure was re-refined using PDB-REDO (Joosten, R.P. et al., 2014). The final data collection and refinement statistics for the MINDY1:K48-Ub2 complex structure is shown in Table 1. All Figs were made using PyMOL (http://pymol.org).

**MINDY2:K48-Ub2**

The MINDY2-catalytic domain C266A mutant construct (residues 241-504) was expressed in *E.coli* BL21(DE3) cells as described. Purified MINDY2 protein in 50 mM Tris-HCl pH 7.5, 150 mM NaCl, and 10 mM DTT was mixed with K48-linked diUb and concentrated to a final concentration of 18.0 mg/ml. The crystals grew grew in hanging drops (24 well plates ) from conditions with 0.05 M Potassium phosphate monobasic and 20% PEG 8000. The crystals were flash frozen in cryoprotectant containing 0.05 M Potassium phosphate monobasic and 5 % PEG 8000 and 30% PEG 8000. Diffraction data were collected at ID29 beamline, ESRF, France (wavelength 0.97625 Å). The data sets were processed, and structures determined as for MINDY1:K48-Ub2 structure.

**P138AC137A:K48-Ub2**

The MINDY1 P138A C137A mutant construct (residues 110-384) was expressed in *E. coli* BL21(DE3) cells and purified as described. Purified protein in 50 mM Tris-HCl pH 7.5, 150 mM NaCl, and 10 mM DTT was mixed with K48-linked diUb and concentrated to a final concentration of 14.0 mg/ml. The crystals were grown in hanging drop 24 well plates against the well solution of 0.2 M sodium malonate pH 7.0 and 20 % PEG 3350. The crystals were flash frozen in cryo-protectant containing 0.2 M sodium malonate pH 7.0 and 35% PEG 400. Diffraction data were collected at ID30B beamline, ESRF, France (wavelength 0.99187 Å).The data sets were processed, and structures determined as for MINDY1:K48-Ub2 structure.

**MINDY2$^{apo}$**

The catalytic domain of MINDY2 (residues 241-504) was expressed in *E.coli* BL21(DE3) cells and purified as described. MINDY2$^{apo}$ protein in 50 mM Tris-HCl pH 7.5, 150 mM NaCl, and 10 mM DTT. The crystals were grown from hanging drops containing an equal volume of protein (12.5 mg/ml) and mother liquor containing 0.1 M Bis Tris pH 6.0, 0.2 M $MgCl_2$ and 25 % PEG 3350. The crystals were flash frozen in cryoprotectant containing 0.1 M Bis Tris pH 6.0, 0.2 M $MgCl_2$ and 35 % PEG 400. Diffraction data were collected at ID29 beamline, ESRF, France (wavelength 1.07252 Å). The data sets were processed using XDS (Kabsch, 2010) and then scaled using AIMLESS (Evans and Murshudov, 2013). The structure of the MINDY2$^{apo}$ was solved by molecular replacement (MoRDa) using MINDY1$^{apo}$ (PDB ID: 5JKN) as search model.

**MINDY1 Y114F**

The catalytic domain of MINDY1 Y114F construct (residues 110-384) was expressed and purified as described. The crystals were grown from hanging drops containing an equal volume of protein (11.5 mg/ml) and mother liquor containing 0.1 M Tris-HCl pH 8.5, 1.5 M Ammonium phosphate dibasic. The crystals were flash frozen in cryoprotectant containing 0.1 M Tris-HCl pH 8.5, 3.4 M Sodium malonate pH 8.0 and 20% glycerol. Diffraction data were collected at ID23-2 beamline, ESRF, France (wavelength 0.87313 Å). The structure of the Y114F was solved by molecular replacement (Phaser) using MINDY1$^{apo}$ (PDB ID: 5JKN) as search model.

**MINDY1 P138A**

MINDY1 P138A (residues 110-384) was expressed and purified as described. The crystals were grown from hanging drops containing an equal volume of protein (11.05 mg/ml) and mother liquor containing 0.1 M Bis Tris Propane pH 7.0 and 0.7 M sodium citrate tribasic dihydrate. The crystals were flash frozen in cryoprotectant containing 0.1 M Bis Tris Propane pH 7.0 and 0.7 M sodium citrate tribasic dihydrate and 30 % PEG 400. Diffraction data were collected at ID23-1 beamline, ESRF, France (wavelength 0.97625 Å). The structure of the P138A was solved by molecular replacement (Phaser) using MINDY1$^{apo}$ (PDB ID: 5JKN) as search model.

**MINDY1 T335D**

MINDY1 T335D (residues 110-384) was expressed and purified as described. The crystals were grown from hanging drops containing an equal volume of protein (12.0 mg/ml) and mother liquor containing 0.1 M Hepes-Na pH 7.5 and 0.8 M Potassium sodium tartrate tetrahydrate. The crystals were flash frozen in cryoprotectant containing 0.07 M Hepes-Na pH 7.5, 0.52 M Potassium sodium tartrate tetrahydrate and 35 % glycerol. Diffraction data were collected at I03 beamline, Diamond, UK (wavelength 0.97628 Å). The structure of the T335D was solved by molecular replacement (Phaser) using MINDY1$^{apo}$ (PDB ID: 5JKN) as search model.

**Sequence conservation**

Analysis of sequence conservation was performed by aligning sequences of MINDY1 from 19 different species: *Homo sapiens* (human), *Gorilla gorilla* (gorilla), *Pan troglodytes* (chimpanzee), *Pongo abelii* (orangutan), *Loxodonta africana* (african elephant), *Bos taurus* (cattle), *Mus musculus* (mouse), *Rattus norvegicus* (rat), *Monodelphis domestica* (opossum), *Lipotes vexillifer* (dolphin), *Felis catus* (cat), *Equus caballus* (horse), *Ovis aries* (sheep), *Ursus maritimus* (polar bear), *Physeter macrocephalus* (sperm whale), *Xenopus tropicalis* (frog), *Takifugu rubripes* (Japanese pufferfish), *Danio rerio* (zebra fish). Consurf analysis was performed using the webserver: https://consurf.tau.ac.il/

**Small angle X-ray scattering (SAXS)**

Synchrotron radiation X-ray scattering patterns from MINDY2-apo, in complex with Ub$^{Prg}$, and polyubiquitin were collected at the EMBL P12 beamline of the storage ring PETRA III (DESY, Hamburg, Germany) (Blanchet et al., 2015). Images were recorded using a photon counting Pilatus-6M detector at a sample to detector distance of 3.0 m and a wavelength (λ) of 0.12 nm covering the range of momentum transfer $0.01 < s < 7\ nm^{-1}$ with $s=4\pi \sin\theta/\lambda$, where 2θ is the scattering angle. To obtain data from monodisperse samples (Mindy2-apo, Mindy2-Ub1$^{prg}$, Mindy2-Ub2, Mindy2-Ub3, Mindy2-Ub4 and Mindy2-Ub5), samples were passed through size exclusion chromatography (Superdex 200 10/300) directly coupled to the SAXS instrument

(SEC-SAXS). Only frames corresponding to the main elution peaks were considered. The SAXS data collected from the buffer components before the elution peak was used for background subtraction. The overall structural parameters of the apo construct derived from the SAXS data are compatible with a monomeric species whilst those for the complexes show trends typical of complex formation and thus confirm the binding of the increasing number of ubiquitin molecules to Mindy2 (Table S3).

One second sample exposures were recorded throughout the entire chromatography step. Buffer S (20 mM HEPES pH 7.5, 100 mM NaCl, 5mM DTT) was used as mobile phase. 100 $\mu$l of purified sample were injected onto a Superdex 200 10/300 (GE Healthcare) column and the flow rate was set to 0.5 ml/min. SAXS data were also recorded from macromolecule-free fractions corresponding to the matched solvent blank. Data reduction to produce final scattering profiles of MINDY 2 constructs was performed using standard methods. Briefly, 2D-to-1D radial averaging was performed by the SASFLOW pipeline (Franke *et al*, 2017) CHROMIXS was used for visualisation and reduction of the SEC-SAXS datasets (Panjkovich & Svergun, 2018). Aided by the integrated prediction algorithms in CHROMIXS the optimal frames within the elution peak and the buffer regions were selected. Single buffer frames were then subtracted from sample frames one by one, scaled and averaged to produce the final subtracted curve. The radius of gyration $R_G$ was computed for each construct by Guinier approximation (Guinier, 1939). The molecular mass (MM) of the solutes was evaluated based on the concentration independent approach using Porod invariant (Porod, 1951) as implemented in the ATSAS package (Hajizadeh *et al*, 2018). The indirect Fourier transform of the SAXS data and the corresponding probable real space pair distance distribution (p(r) versus r profile) of the MINDY2 constructs were calculated using GNOM (Svergun, 1992) yielding also the particle diameter $D_{max}$. The theoretical curves were calculated from the atomic models with CRYSOL (Svergun *et al*, 1995). In case of systematic deviations, normal mode analysis as implemented in SREFLEX (Panjkovich & Svergun, 2016) was employed to refine the crystallographic models. Possibilities of having mixtures of complexes with distinguishable Ub2 binding positions were analysed with OLIGOMER (Konarev et al., 2003).

The SAXS data (as summarized in **Table S3**) as well as fits to the curves computed from the crystal structures and from the refined models have been deposited into the Small-Angle Scattering Biological Data Bank (SASBDB) (Valentini *et al*, 2015).
